# Innovative mutant screening identifies *TRANSPARENT TESTA7* as a player in seed oil/protein partitioning

**DOI:** 10.1101/2024.11.18.624101

**Authors:** Alain Lécureuil, Massimiliano Corso, Stéphanie Boutet, Sophie Le Gall, Regina Niñoles, Jose Gadea, Philippe Guerche, Sophie Jasinski

## Abstract

Brassicaceae species mainly accumulate oil and protein in their seeds, which are essential to human life as a source of food, but also as animal feed and resources for green chemistry. To date, Brassicaceae crops such as rapeseed have been selected mainly for their oil content. However, there is a growing interest in their seed protein content. A strong negative correlation between oil and protein content makes it difficult to increase both compounds simultaneously. In this study, an *Arabidopsis thaliana* homozygous EMS mutant library was screened by near-infrared spectroscopy for seed oil and protein content, with the aim of identifying mutants with impaired oil-protein correlation. The mutant most affected in this correlation was found to be in the *TRANSPARENT TESTA7* gene, which is involved in the flavonoid biosynthetic pathway. Analysis of different mutants in the flavonoid pathway revealed that the *tt7* mutants were the only ones to show such a significant reduction in seed oil content, highlighting a phenotype never described before for the *tt7* mutants and suggesting a specific role for TT7 in the interplay between the oil and flavonoid biosynthetic pathways. Untargeted metabolomic analysis allowed the identification of metabolic features that are highly accumulated and specific to *tt7* seeds compared to the other genotypes and genetic analysis established that the accumulation of kaempferol-3-O-rhamnoside seems to be responsible for the seed oil reduction of *tt7* mutants.

**Significance Statement:** Brassicaceae species accumulate oil and protein in their seeds and understanding how the partitioning of these compounds is regulated is necessary to engineer seeds for specific purposes. By screening an Arabidopsis EMS mutant library, we identified mutants affected in seed oil/protein partitioning, including *tt7*, highlighting a link between oil and flavonoid biosynthetic pathways, that we explore further in this paper.

## INTRODUCTION

Seed storage compounds consist mainly of starch, proteins and oil and are of critical importance for human food, feed and industrial uses. In oleo-proteaginous species such as rapeseed, seed oil has been the main qualitative determinant of the economic value of the harvested seed. Today, however, there is increasing interest in the seed protein content of crop species. Indeed, global population growth and rising standards of living will lead to dietary changes that will increase human consumption of plant-based oil and protein as well as an increased demand for oilseed meals for animal feed. Rapeseed is one of the major oil crops grown on many continents and contributes more than 15% of the world’s edible oil supply. It is the most important oleo-proteaginous crop grown in Western Europe and is indeed a good candidate for increasing plant protein production, which would enable this continent to become less dependent on imported plant proteins. Of course, this should be achieved without compromising seed yield and oil content, and taking into consideration oil and protein quality.

The biosynthetic pathways of seed storage proteins and oil are well described and genes encoding the key enzymes of these pathways have been identified in several oleo-proteaginous species (Shewry *et al*., 1995; Ohlrogge and Jaworski, 1997; Baud *et al*., 2008; Baud and Lepiniec, 2010). However, the genes and mechanisms that determine the differential partitioning of seed reserves into the major storage components remain largely unknown. Identification of these factors will be necessary if we are to succeed in designing high-yielding crops that produce ‘customised’ seeds with specific nutritional value by regulating not only the amount but also the distribution of storage compounds. A strong negative correlation between seed oil and protein content has been observed in oil-storing seeds such as rapeseed (Grami *et al*., 1977; Jolivet *et al*., 2013), sunflower (Li *et al*., 2017), soybean (Kambhampati *et al*., 2020) or the model plant *Arabidopsis thaliana* (Jasinski *et al*., 2018), suggesting that seed filling is highly constrained in these species and that manipulation of both components independently may not be obvious. Several QTL/GWAS studies in these species support this hypothesis, as oil and protein content QTL are often co-localised but have opposite effects (Chung *et al*., 2003; Nichols *et al*., 2006; Bouchet *et al*., 2014; Hwang *et al*., 2014; Jasinski *et al*., 2016). Accordingly, attempts to separate oil and protein content QTL in soybean have been unsuccessful (Chung *et al*., 2003; Nichols *et al*., 2006), reinforcing the hypothesis that the same genes control both traits. However, some results are not fully consistent with this assertion. In soybean, Hwang *et al*. (2014) identified a SNP associated with both higher protein and oil content, and QTL specific for oil or protein content have been identified in both rapeseed and Arabidopsis (Bouchet *et al*., 2014; Jasinski *et al*., 2016). These QTL are of great interest for improving both traits independently. Furthermore, mutant studies in Arabidopsis have shown that a decrease in the amount of one storage compound does not necessarily lead to a compensatory increase in the other compounds (Finkelstein and Somerville, 1990; Focks and Benning, 1998). This suggests that protein and oil content may be uncoupled.

It is therefore of interest to unravel the existing correlation between seed oil and protein accumulation and to learn how to break or at least weaken this correlation to manage both components independently. We aim to identify genetic factors controlling seed oil and protein relative accumulation in oleo-proteaginous species and to perform functional analysis of these genes. This should help understanding this negative correlation and thus provide the tools to manipulate the oil/protein ratio. An efficient way to identify genes involved in a biological process is to carry forward genetic approaches such as QTL mapping, genome wide association studies (GWAS) or screening of mutant libraries for example. Over the last decade, whole-genome sequencing has proved to be a very powerful and rapid tool for identifying mutations induced by mutagens in Arabidopsis as well as in several crops (Candela *et al*., 2015; Thole and Strader, 2015; Tang *et al*., 2020). Thus, to tackle the genetic regulation of seed oil and protein correlation in *Arabidopsis thaliana*, an innovative strategy was adopted. First, a homozygous EMS-mutant library was screened by near-infrared spectroscopy (NIRS) to determine the seed oil and protein content of the approximately 600 lines in the library. Screening of a homozygous (as opposed to heterozygous) mutant collection is essential to allow phenotyping of homogeneous seed pools. Second, the Delta Seed Composition (DSC) trait defined by Marmagne *et al*. (2020) for seed carbon and nitrogen content, was extended to seed oil and protein content. DSC represents the residuals of the orthogonal regression of protein content on oil content. It is highly heritable and is an indicator of the closeness of the relationship between the two compounds for a given genotype (Marmagne *et al*., 2020). This allowed us to identify genotypes that produce seeds with oil and protein contents that do not follow the usual negative correlation. In this study, the mutant with the highest DSC (*HEM_115*) was studied in more detail. A mapping-by-sequencing approach followed by insertion mutant analysis and functional complementation demonstrated that a mutation in *TRANSPARENT TESTA7* (*TT7*), encoding a protein involved in the biosynthesis of flavonoids, is responsible for the high DSC of the *HEM_115* line. In this paper, we studied *tt7* and other mutants of the flavonoid pathway, which allowed us to gain new insight into the interplay between flavonoid and oil biosynthetic pathways.

## RESULTS

### 1. Near-Infrared spectroscopy is suitable for screening mutants with impaired oil/protein content

In order to identify mutants affected in the composition of seed storage compounds, we took advantage of a library of homozygous EMS mutant (HEM) lines developed from the Col-0 accession by Capilla-Perez *et al*. (2018). This mutant collection was used in a forward genetic screen targeting seed filling phenotypes. For this purpose, seed stocks of 591 lines from the biological resource centre (BRC) were directly phenotyped by NIRS and their seed oil and protein contents were estimated using NIRS models that were previously developed (Jasinski *et al*., 2016) (Figure 1a). This collection showed a wide range of variation for seed oil content from 19.5% to 45.0%, with a mean of 35.1% and a standard deviation (SD) of 3.4% (Figure 1a and Table 1). Similarly, a wide range of seed protein content was observed from 15.0% to 24.7% with a mean of 19.7% and a SD of 1.5% (Figure 1a and Table 1). A strong negative correlation between oil (O) and protein (P) contents was observed, as already described for Arabidopsis (Jasinski *et al*., 2018) and other plant species that mainly accumulate oil in their seeds, such as rapeseed (Grami *et al*., 1977; Boulard *et al*., 2015; Jolivet *et al*., 2013) or sunflower (Li *et al*., 2017), but also in species that mainly accumulate protein, such as soybean (Chung *et al*., 2003). As expected, most genotypes showed seed oil and protein contents following the P/O regression line, i.e. close to a null DSC (Figure 1a). However, some genotypes displayed a high DSC (positive or negative), indicating that the negative correlation was removed or relaxed in these genotypes. Calculation of DSC values showed that they varied from –9.7 to 4.7 with a mean of 0.0 and a SD of 1.7 in the HEM collection (Table 1). Mutant lines with high DSC, either negative or positive, as well as lines with contrasted seed oil and protein contents were considered for our study. In total, nine mutants were selected for further analysis (Figure 1a).

**Figure 1.**
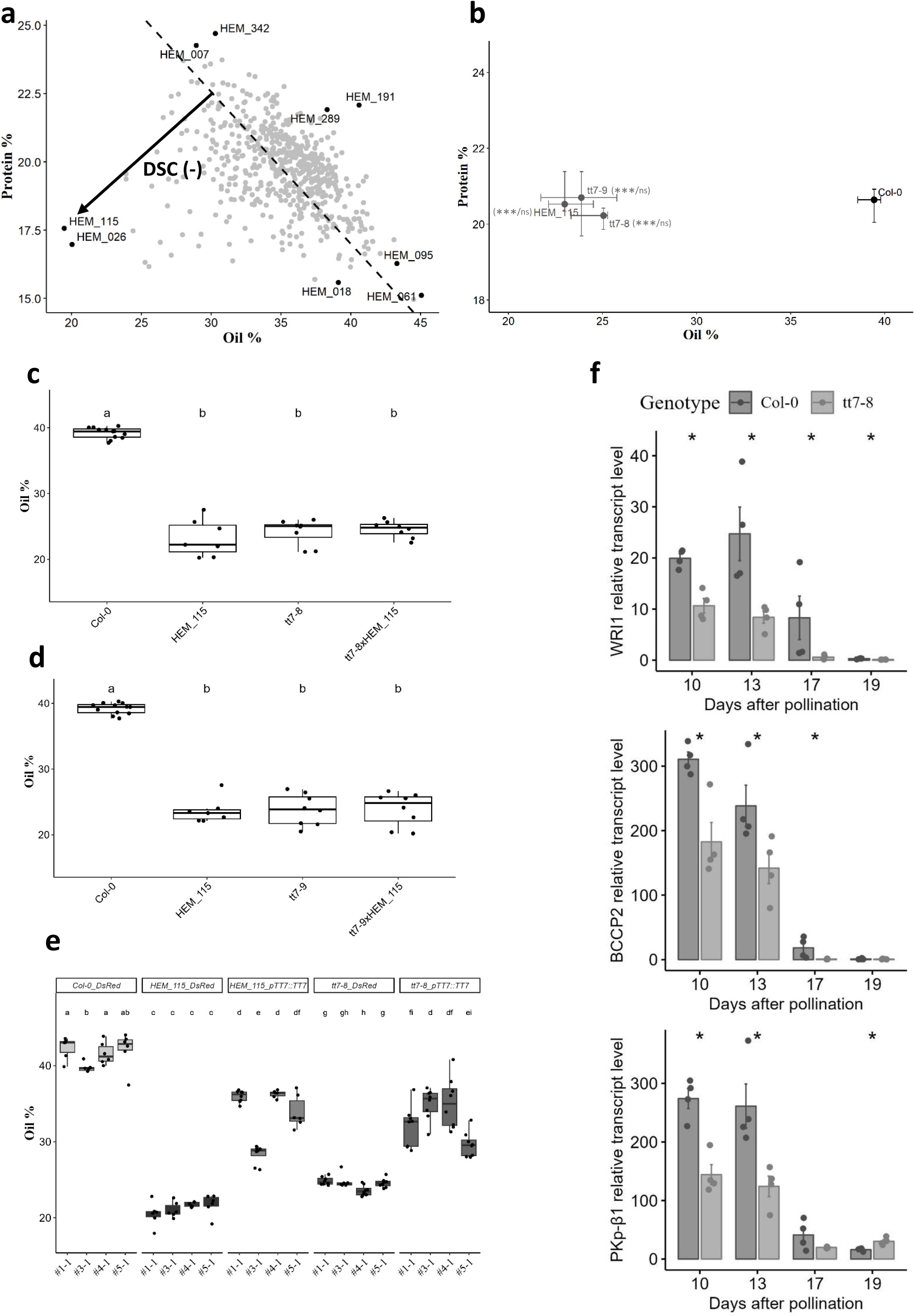
mutation in *TT7* is responsible for the seed composition phenotype of the *HEM_115* mutant. (a) Seed oil and protein contents of 591 HEM lines. Each dot corresponds to a mutant line. Dots in black are the nine mutants studied further with their corresponding name indicated (*HEM_*). DSC (Delta Seed Composition), which corresponds to the orthogonal distance between one point and the P/O regression line (dashed line) is indicated by an arrow. The DSC is negative when it is below the regression line and positive when above the regression line. (b) Median of seed oil and protein contents are represented for Col-0, *HEM_115*, *tt7-8* and *tt7-9*. Bars show the 25–75% quartiles (n ≥ 8). Col-0 is in black, the other genotypes are in grey. A Kruskal-Wallis statistical test, followed by a post-hoc Man & Whitney test were performed to compare each mutant to Col-0 for both traits. Significance of comparison is indicated in bracket (oil/Protein), ***P < 0.001, ns=not significant. (c, d) Median of seed oil content is represented for Col-0 and the three classes of genotypes from a cross between *HEM_115* and *tt7-8*: *HEM_115*, *tt7-8* and *HEM_115 x tt7-8* (c) or a cross between *HEM_115* and *tt7-9*: *HEM_115*, *tt7-9* and *HEM_115 x tt7-9*. Boxes show the 25–75% quartiles, the median value (inner horizontal line), and whiskers extending from the lower to the upper adjacent values. A Kruskal-Wallis statistical test, followed by a post-hoc pairwise Wilcoxon test were performed to compare all pairs of genotypes. Genotypes with the same letter are not significantly different from each other, while genotypes with different letters are significantly different (P ≤ 0.05). (e) Median of seed oil content for Col-0 DsRed, *HEM_115* DsRed, *tt7-8* DsRed and Col-0, *HEM_115* and *tt7-8* transformed with *pTT7::TT7* DsRed lines. For each construct, T2 or T3 seeds from four independent homozygous lines were subjected to NIRS. A Kruskal-Wallis statistical test, followed by a post-hoc pairwise Wilcoxon test were performed to compare all pairs of genotypes. Genotypes with the same letter are not significantly different from each other, while genotypes with different letters are significantly different (P ≤ 0.05). (f) Expression levels of *WRI1*, *BCCP2* and *PKp-β1* relative to *EF1a* and *PP2AA3* reference genes in *tt7-8* and wildtype during seed development. The values of the bars are means of four biological replicates, indicated by points. Each point corresponds to the mean of two technical repeats. For each day, *tt7-8* was compared to Col-0 with a Man & Whitney statistical test. Significance of comparison is indicated, ***P < 0.001, **P < 0.01 and *P < 0.05.

**Table 1:** statistics for seed oil and protein contents of the HEM lines.

|  | Mean | Range | SD |
| --- | --- | --- | --- |
| **Oil** | 35.1 | 19.5 - 45.0 | 3.4 |
| **Protein** | 19.7 | 15.0 - 24.7 | 1.5 |
| **DSC** | 0.0 | -9.7 - 4.7 | 1.7 |

First, we aimed to confirm the phenotype of each mutant on new seed batches, as the NIRS screening was carried out on seeds supplied directly by the BRC. To do this, the nine mutants and the wildtype Col-0 (10 replicates per genotype) were grown in parallel in randomised blocks in the greenhouse, with trays moved every two days to homogenise the conditions. Mature seeds of all the plants were subjected to NIRS to estimate seed oil and protein contents. Although the absolute values of seed oil and protein contents were not similar between this experiment and the screening experiment, the position of each genotype on the P/O graph was well conserved, showing that the seed phenotype of each mutant was indeed genetically controlled and reproducible (Figure S1a). For line *HEM_026*, only one out of ten plants produced seeds and this line was therefore not studied further. This second phenotyping allowed us to determine which mutants were significantly different from wildtype Col-0 for at least one trait between seed oil and protein content. The mutant *HEM_095* was not significantly different from the wildtype for either oil or protein content (Figure S1a) and was therefore not studied further.

In addition, both seed oil and protein contents were measured in six out of the nine mutants and the wildtype by reference methods, gas chromatography and phenolic extraction, respectively. As shown in Figure S1b, c, for both traits, NIRS and corresponding wet-chemistry method values were highly significantly correlated, but with a higher correlation coefficient for oil content (R=0.99) than for protein content (R=0.61). These results demonstrated that NIRS is suitable for the screening of mutants with an impaired oil/protein correlation.

### 2. *TRANSPARENT TESTA 7* plays a role in seed oil filling

The *HEM_115* mutant displayed the highest DSC (*HEM_026* excluded) from the HEM library. It showed a strongly impaired oil/protein correlation, with a very low oil content related to its protein content. In order to identify the causal gene of this phenotype, a mapping-by-sequencing approach was carried out. For this purpose, an F2 population of 150 individuals from a cross between Col-0 and *HEM_115* was grown in parallel with the parents and their F3 seeds were subjected to NIRS analysis (Figure S1d). The F3 seed pools displayed a wide range of variability from mutant to wildtype phenotype, with approximately 1/4 of these F3 pools displaying a mutant phenotype, as expected for segregation of a single mutated locus. Twenty-four F2 plants, which produced F3 seeds with a mutant phenotype and were therefore expected to carry the mutation of interest at the homozygous state, were selected for deep sequencing in pool. Sequence analysis allowed the identification of a candidate region of approximately 5 Mb on the top of chromosome 5 from 0 to 5 Mb. Detailed analysis of the mutations in this region led to nine candidate genes (Table 2). Among these nine candidates, seven had a mutation leading to a non-synonymous amino-acid modification, one had a mutation in the splice-site region and one, *At5g07990* (*TRANSPARENT TESTA 7*, *TT7*), displayed a premature STOP codon, leading to a truncated protein of 39 amino-acids in *HEM_115* instead of a 513 amino-acid protein in the wildtype (Figure S1e). *TT7* encodes a cytochrome P450 75B1 monooxygenase involved in flavonoid biosynthesis (Abrahams *et al*., 2002; Appelhagen *et al*., 2014, see also below and Figure 4a) and other mutants in the same pathway, in particular *tt2* (Chen *et al*., 2012; Wang *et al*., 2014), *tt4* (Xuan *et al*., 2018), *tt8* (Chen *et al*., 2014) and *ttg1* (Chen *et al*., 2015) have been reported to show altered seed oil content. Taken together, these results highlighted *TT7* as the best candidate responsible for the *HEM_115* phenotype.

**Table 2:**
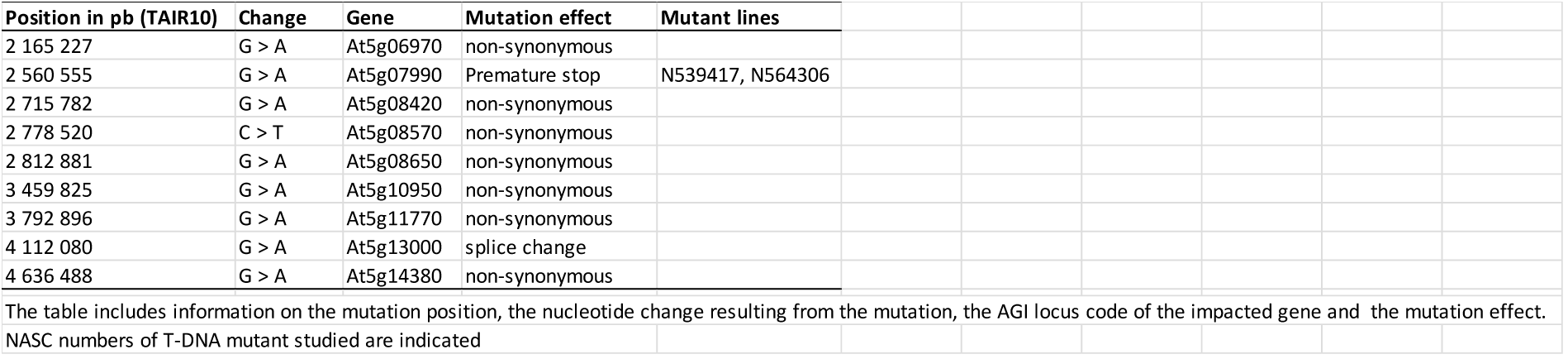
List of the mutations identified as candidate causal mutations in the HEM_115 mutant.

To determine whether the mutation in *TT7* was indeed responsible for the seed filling phenotype of *HEM_115*, two T-DNA mutants in *TT7*, *tt7-8* and *tt7-9* (see M&M) were analysed. Molecular characterization of the mutants revealed that *tt7-8* and *tt7-9* carried a T-DNA insertion combined with a 34 or 35-base pair deletion in the second exon of the *TT7* gene respectively. Similar to *HEM_115*, both *tt7* alleles displayed a reduced seed oil content compared to Col-0 (with a median of 25.0%, 23.9%, 23.0% and 39.4% for *tt7-8*, *tt7-9*, *HEM_115* and Col-0 respectively) with no compensation in protein content (Figure 1b). This decrease in seed oil content was accompanied by an altered seed oil composition in all three alleles (Figure S1f). This result suggested that the reduced seed oil content of *HEM_115* was due to the mutation in the *TT7* gene and demonstrated that *TT7* is involved in seed oil accumulation. To ensure that the phenotype of *HEM_115* was not due to other mutations in any of the other eight candidate genes, allelism tests were carried out. *HEM_115* was crossed with *tt7-8* and *tt7-9* and forty F2 seeds per cross were sown, plantlets were genotyped and F3 seeds from each plant were phenotyped by NIRS. For both crosses, F3 seed lots from F2 heterozygous plants (*HEM_115*/+ *tt7-8*/+ or *HEM_115*/+ *tt7-9*/+) displayed reduced seed oil content compared to Col-0, like each single mutant (Figure 1c, d), demonstrating that the mutation in the *TT7* gene is indeed responsible for the *HEM_115* phenotype. To further validate the role of *TT7* in seed oil accumulation, *HEM_115* and *tt7-8* mutants were transformed with a construct harbouring the WT *TT7* genomic sequence fused to a 2kb-*TT7* promoter and a DsRed fluorescent marker gene (see M&M). Homozygous T2 plants from T1 harbouring a single T-DNA insertion locus were grown and their T3 seeds subjected to NIRS (Figure 1e). As controls, Col-0, *HEM_115* and *tt7-8* plants transformed with DsRed alone were analysed. As shown in Figure 1e, *HEM_115_DsRed* and *tt7-8_DsRed* plants displayed a reduced seed oil content compared with Col-0_DsRed plants, whereas *HEM_115* and *tt7-8* plants transformed with p*TT7::TT7* displayed a rescued phenotype.

Taken together, these results demonstrated that *TT7* plays a role in seed oil accumulation by positively affecting seed oil content. To further explore the mechanism by which seed oil content was affected in *tt7* mutants, we examined the transcription levels of *WRINKLED1* (*WRI1*), a master transcription factor in the regulation of plant oil biosynthesis (Focks and Benning, 1998; Baud *et al*., 2007; Cernac and Benning, 2004), and two of its target genes: *PLASTIDIAL PYRUVATE KINASE β1*(*PKp-β1*) and *BIOTIN CARBOXYL CARRIER PROTEIN 2*(*BCCP2*) (Baud *et al*., 2009; Maeo *et al*., 2009) in WT and *tt7-8*. A quantitative RT-PCR approach was performed on seeds at different developmental stages from 10 days after pollination (DAP) to mature seeds, as the transcript level of these genes is maximal in this time window (Baud *et al*., 2009). The qRT-PCR analysis showed that the transcript level of the three genes was significantly reduced in *tt7-8* compared to WT at almost all developmental stages analysed (Figure 1f). This result suggested that the reduction in seed oil content observed in *tt7-8* may be due, at least in part, to a reduction in *WRI1* expression and consequently a reduction in the expression of its target genes.

### 3. The reduced seed oil content of *tt7* is controlled by the seed coat

Seed oil synthesis occurs after embryo morphogenesis during the seed maturation phase from 7 to 20 DAP and the embryo accumulates 90% of the seed oil (Baud and Lepiniec, 2009). To understand why *tt7* mutants displayed reduced seed oil content, their embryo development was compared with that of Col-0 (Figure 2a). As described before for *tt7-7* (Niñoles *et al*., 2023), the three *tt7* mutants analysed showed exactly the same kinetics of embryo development as Col-0, namely the embryo is at globular stage 4 DAP, at early heart stage 6 DAP, at torpedo stage 8 DAP, at upturned-U stage 10 DAP and at mature cotyledon stage 12 DAP. Nevertheless, we noticed that the seed size of the three *tt7* mutants was reduced compared with Col-0 (Figure 2b), a phenotype already observed for *tt7-1* (Debeaujon *et al*., 2000; Lu *et al*., 2008). Since oil content is expressed as a percentage of seed dry mass, the observed reduced seed oil content cannot be explained by the reduced seed size/mass. This showed that the mutation in the *TT7* gene did not impact embryo development and therefore the reduced seed oil content observed in *tt7* mutants was not a consequence of an impaired embryo development.

**Figure 2.**
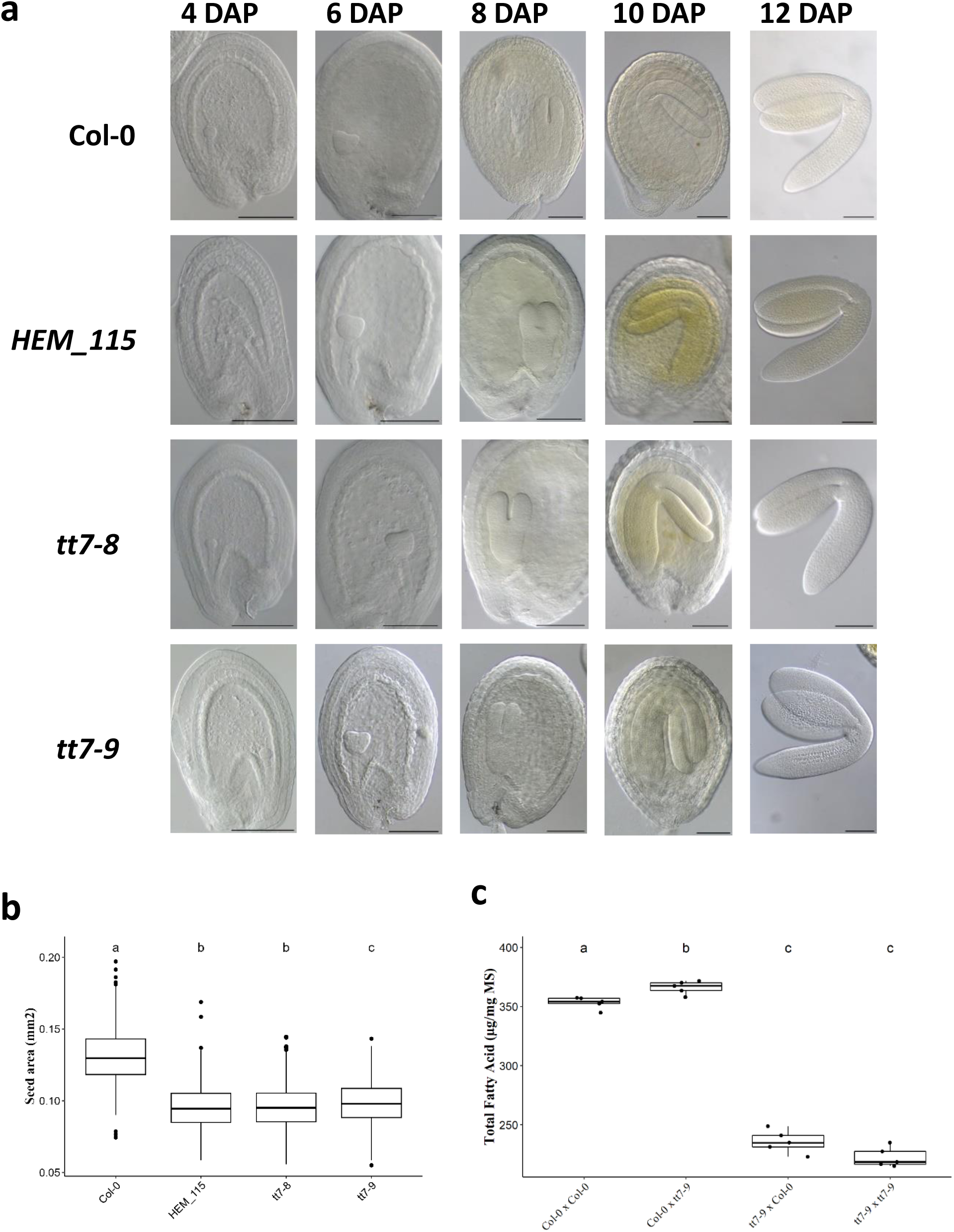
*tt7* mutants have an embryo development similar to wildtype. (a) Developing seeds of Col-0, *HEM_115*, *tt7-8* and *tt7-9* mutants were cleared, and embryos were observed at 4, 6, 8,10 and 12 days after pollination (DAP). Scale bars = 100 µm. (b) Seed area of Col-0, *HEM_115*, *tt7-8* and *tt7-9*. Boxes show the 25–75% quartiles, the median value (inner horizontal line), and whiskers extending from the lower to the upper adjacent values. At least 440 seeds have been measured for each genotype. Points indicate outliers’ values. A Kruskal-Wallis statistical test, followed by a post-hoc pairwise Wilcoxon test were performed to compare all pairs of genotypes. Genotypes with the same letter are not significantly different from each other, while genotypes with different letters are significantly different (P ≤ 0.05). (c) Total fatty acid content of F1 seeds from the four indicated crosses. Boxes show the 25–75% quartiles, the median value (inner horizontal line), and whiskers extending from the lower to the upper adjacent values. A Kruskal-Wallis statistical test, followed by a post-hoc pairwise Wilcoxon test were performed to compare all pairs of genotypes. Genotypes with the same letter are not significantly different from each other, while genotypes with different letters are significantly different (P ≤ 0.05).

*TT7* is mainly expressed in the seed coat, or testa (Belmonte *et al*., 2013), a maternal tissue, where it is involved in flavonoid biosynthesis (Shirley *et al*., 1995). However, seed oil synthesis mainly takes place in the embryo. To determine whether *TT7* acts maternally or zygotically in affecting oil biosynthesis, we made reciprocal crosses between the wildtype Col-0 and the *tt7-9* mutant and we measured the seed oil content in the respective F1 seeds by gas chromatography (M&M, Figure 2c). As expected, F1 seeds derived from *tt7-9* x *tt7-9* crosses showed a reduced oil content compared to F1 seeds derived from Col-0 x Col-0 crosses. The difference was of the same order as that observed for the *tt7-9* mutant compared to the Col-0. F1 seeds derived from Col-0 x *tt7-9* crosses (with Col-0 testa) showed the same oil content as F1 seeds from Col-0 x Col-0 crosses. In contrast, F1 seeds from *tt7-9* x Col-0 crosses (with mutant *tt7* testa) showed the same oil content as the F1 seeds from *tt7-9* x *tt7-9* crosses. In these two types of crosses (Col-0 x *tt7-9* and *tt7-9* x Col-0), the embryos had the same heterozygous genotype, whereas the testa, derived from maternal tissue, had either the Col-0 or *tt7-9* genotype, demonstrating that the genotype of the testa determines the oil content of the embryo. These results, together with the fact that *tt7* embryos showed normal morphology and development, indicated that the reduced seed oil content observed in *tt7* mutants is maternally inherited, through the seed coat.

### 4. *TT7* is involved in seed coat differentiation

In wildtype Arabidopsis, proanthocyanidins (PAs, also called condensed tannins) accumulate in the endothelium, the innermost cell layer of the inner integument. During seed desiccation, these tannins are oxidised, leading to the brown colour of wildtype Arabidopsis seeds. As already published for many *tt7* alleles (Shirley *et al*., 1995; Koornneef *et al*., 1982; Debeaujon *et al*., 2000; Abrahams *et al*., 2002), the three *tt7* alleles studied in this paper exhibited pale brown seeds (Figure 3a) compared to brown wildtype seeds, indicating a defect in PA synthesis.

**Figure 3.**
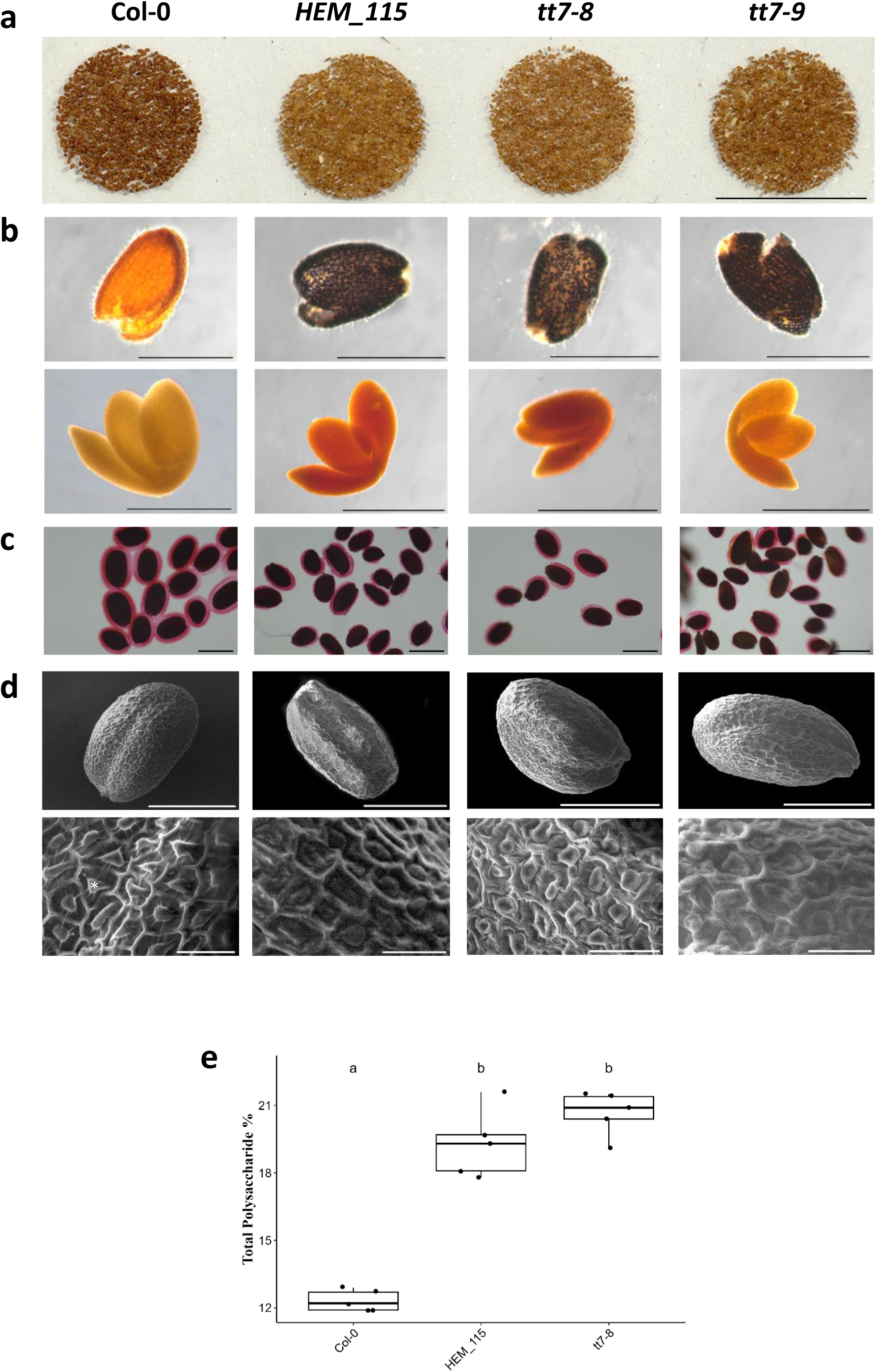
TT7 is involved in seed coat differentiation. (a) Photographs of mature seeds of Col-0, *HEM_115*, *tt7-8* and *tt7-9*. Scale bars = 1 cm. (b) Mature seeds of Col-0, *HEM_115*, *tt7-8* and *tt7-9* were dissected and seed coats and embryos were stained separately for starch with Lugol solution and observed under binocular microscope. Scale bars = 500 µm (c) Col-0, *HEM_115*, *tt7-8* and *tt7-9* seeds were gently mixed in water and ruthenium red (RR) was used to stain adherent pectin (pink). Scale bars = 500 µm. (d) Scanning electron micrographs of whole seed and seed coat details of Col-0, *HEM_115*, *tt7-8* and *tt7-9*. Asterisk indicates a columella and white arrowheads the outer edges of a radial cell wall. Scale bars = 300 µm for whole seeds and 50 µm for details. (e) Box plots of total polysaccharide content. Boxes show the 25–75% quartiles, the median value (inner horizontal line), and whiskers extending from the lower to the upper adjacent values. About 100 mg of seeds from 5 independent plants per genotype were analysed. A Kruskal-Wallis statistical test, followed by a post-hoc pairwise Wilcoxon test were performed to compare all pairs of genotypes. Genotypes with the same letter are not significantly different from each other, while genotypes with different letters are significantly different (P ≤ 0.05).

**Figure 4.**
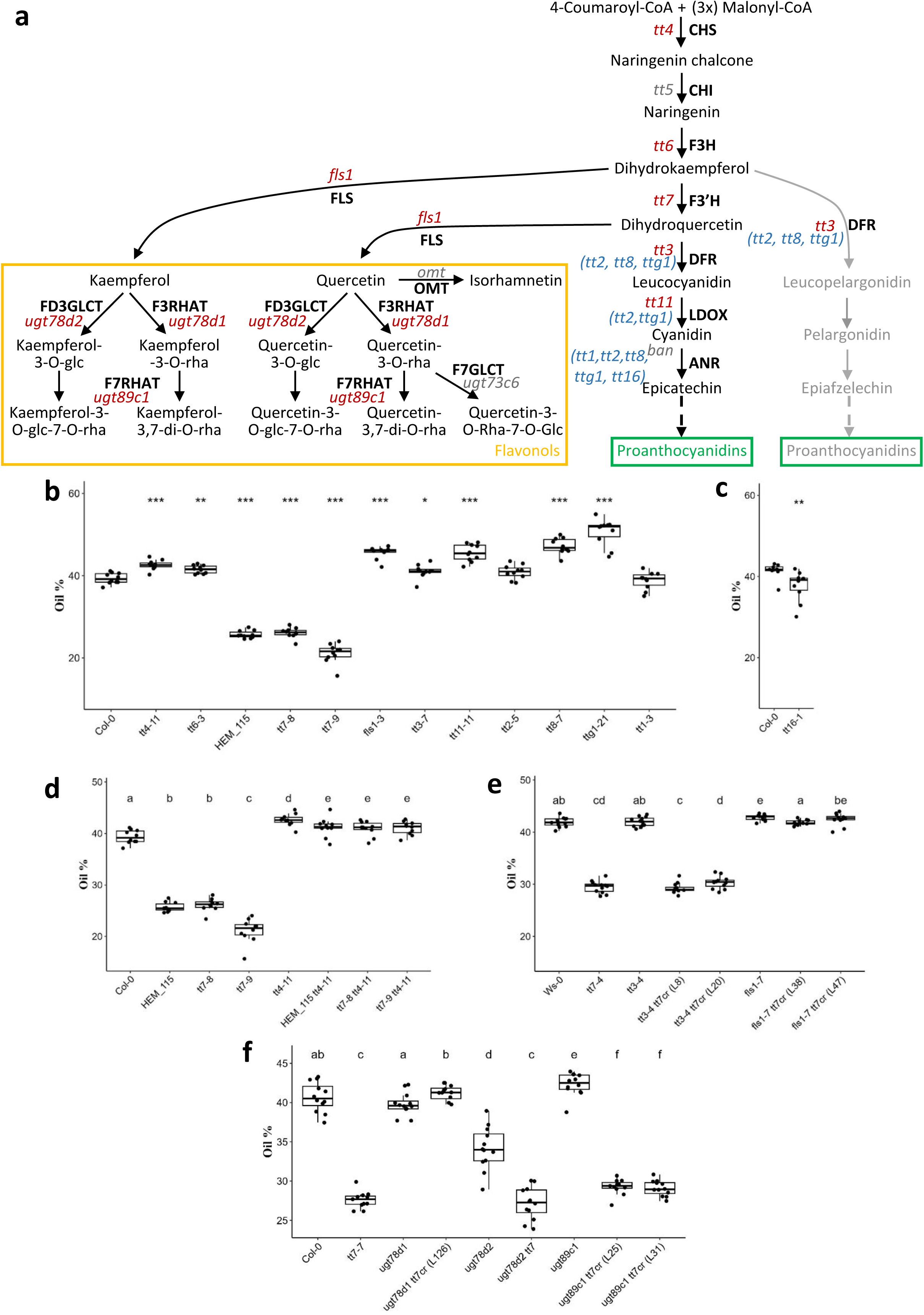
A specific role for TT7 in seed oil content, driven by kaempferol accumulation. (a) Scheme of the flavonoid biosynthetic pathway in Arabidopsis seed. Enzymes are indicated in black capital letters. Mutants for enzymatic steps are indicated in lowercase italic letters, red for the ones studied in this paper, grey for the others. Mutants for regulatory genes are indicated in blue and in bracket beside their target genes. Dashed arrows indicate multiple steps. CHS, chalcone synthase; CHI, chalcone isomerase; F3H, flavanone 3-hydroxylase; FLS, flavonol synthase; F3’H, Flavone 3′-hydroxylase; DFR, dihydroflavonol reductase; LDOX/ANS, leucocyanidin dioxygenase/ anthocyanin synthase; ANR, Anthocyanin reductase; F3RHAT, flavonol-3-O-rhamnosyltransferase; FD3GLCT, flavonol-3-O glucosyltransferase; F7RHAT, flavonol-7-O-rhamnosyltransferase; F7GLCT, flavonol-7-O-glucosyltransferase. (b, c) Median seed oil content of mutants involved in the biosynthesis of flavonoids. Boxes show the 25–75% quartiles, the median value (inner horizontal line), and whiskers extending from the lower to the upper adjacent values. A Kruskal-Wallis statistical test, followed by a post-hoc Man & Whitney test were performed to compare each mutant to Col-0. Significance of comparison is indicated, ***P < 0.001, **P < 0.01 and *P < 0.05, ns=not significant. (d-f) Median seed oil content of simple and double mutants. A Kruskal-Wallis statistical test, followed by a post-hoc pairwise Wilcoxon test were performed to compare all pairs of genotypes. Genotypes with the same letter are not significantly different from each other, while genotypes with different letters are significantly different (P ≤ 0.05).

In addition to oil and proteins, carbohydrates (saccharides and polysaccharides) are the third component found in mature Arabidopsis seeds. In particular, polysaccharides are present in the cell wall and in the mucilage, as Arabidopsis is a myxospermic species. Since *tt7* mutants showed reduced seed oil content (as a percentage of dry mass) without any compensation in protein content (Figure 1b), it was therefore hypothesised that *tt7* mutants would contain more carbohydrates in their seeds. To test this hypothesis, the total cell wall polysaccharide (CWP) content of dried mature seeds of *tt7*-8 and *HEM_115* was measured. As shown in Figure 3e, both *tt7* alleles displayed a strongly increased CWP content compared to Col-0 (19.3 %, 20.9% and 12.2% for *HEM_115*, *tt7-8* and Col-0 respectively). Analysis of the osidic composition of the CWP showed that all the saccharides were increased in the *tt7* alleles compared to Col-0 (Figure S2), with glucose being the most increased (2.3 and 2.1 times in *tt7-8* and *HEM_115*, respectively, compared to about 1.4 for the other saccharides in both *tt7* alleles). This result demonstrated that the *tt7* alleles displayed an overall higher CWP content compared to wildtype.

To gain more insight into which polysaccharides were affected, starch and mucilage were examined in more detail in the three *tt7* mutants. In Arabidopsis, starch is transiently accumulated in the seed coat and embryo during seed development and is almost undetectable in mature dry seeds (Focks and Benning, 1998; Baud *et al*., 2002; Andriotis *et al*., 2010). To determine whether starch accumulation was impaired in the three *tt7* mutants studied in this paper, as it was shown for *tt7-4* and *tt7-7* (Niñoles *et al*., 2023), mature dry seeds were dissected and the seed coat and embryo were separately subjected to Lugol’s staining. As expected, no staining was found in the Col-0 seed coat or in the embryo of mature dry seeds. On the contrary, dark staining was observed in the seed coat of the three *tt7* mutants (Figure 3b), demonstrating that the starch degradation was specifically impaired in the *tt7* seed coat. This is consistent with the high increase in glucose from CWP (Figure S2). In Arabidopsis, mucilage, is produced and deposited in the outer cell layer of the seed coat. Arabidopsis mucilage consists of two layers: a water-extractable outer layer and an adherent inner layer (Macquet *et al*., 2007). The adherent mucilage of the three *tt7* mutants and col-0 was analysed by ruthenium red staining. A pink halo of mucilage was visible around all the seeds of Col-0, whereas relatively few seeds of the three *tt7* mutants had mucilage halos (Figure 3c). Furthermore, when present, the mucilage halo observed around individual *tt7* seeds did not cover the entire seed and was heterogeneous from seed to seed. Similar results were observed for *tt7-4* (Niñoles *et al*., 2023). As mucilage is extruded from the cells of the epidermal layer, scanning electron microscopy was used to examine these cells. As shown in Figure 3d, Col-0 seeds displayed an epidermal layer of hexagonal cells with radial cell walls and a central, volcano-shaped structure known as the columella, whereas *tt7* mutants displayed collapsed columella compared to Col-0, as observed for *tt7-4* (Niñoles *et al*., 2023). This result demonstrated a defect in the differentiation of mucilage-producing cells.

Taken together, these results provide further insight into the description of the *tt7* seed coat developmental defects.

### 5. Interplay between flavonoid and seed oil content: a specific role for *TT7*

The Arabidopsis seed coat accumulates flavonoids (only flavonols and PAs) whose biosynthetic pathway is driven by many genes of the *TRANSPARENT TESTA* (*TT*) family, which encode either transcription factors or enzymes (Chen *et al*., 2023; Appelhagen *et al*., 2014) (Figure 4a). In Arabidopsis, mutants affected in some *TT* genes, such as *tt4*, *tt2*, *tt8*, and *ttg1* have been shown to contain higher seed oil content compared to the corresponding wildtype (Li *et al*., 2018; Xuan *et al*., 2018; Chen *et al*., 2015; Chen *et al*., 2012; Chen *et al*., 2014). However, the *tt16* mutant displayed a reduced seed oil content (Lu *et al*., 2021), suggesting a complex interaction between flavonoid and oil synthetic pathways. To further addressed this interaction, oil content was measured in Arabidopsis knockout mutants affecting 11 genes (6 structural genes, including *TT7*, and 5 regulatory genes) involved in the flavonoid pathway (Figure 4). *TT4* gene encodes the first enzyme of the flavonoid biosynthetic pathway (the chalcone synthase, CHS), hence, the corresponding mutants do not synthetize the major flavonoid compounds (Shirley *et al*., 1995; Bowerman *et al*., 2012; Schulz *et al*., 2016). *tt6* is impaired in the gene encoding the flavanone 3-hydroxylase and accumulates reduced flavonol levels (Shirley *et al*., 1995; Schulz *et al*., 2016). Seeds containing mutations in the *FLAVONOL SYNTHASE* (*FLS*) gene have no or reduced flavonol content and increased PA content (Routaboul *et al*., 2006; Bowerman *et al*., 2012; Niñoles *et al*., 2023). *TT3* encodes a dihydroflavonol-4-reductase and *tt3* mutants lack PAs but accumulate the flavonols quercetin and kaempferol (Shirley *et al*., 1995; Routaboul *et al*., 2006; Schulz *et al*., 2016). *TT11* gene encodes a leucocyanidin dioxygenase (LDOX), also called anthocyanidin synthase (ANS) and *tt11* seeds contain a wildtype flavonol content but no PAs (Appelhagen *et al*., 2011; Bowerman *et al*., 2012). *tt7* seeds do not contain quercetin nor isorhamnetin-derived flavonols, but accumulate large amounts of kaempferol-derived flavonols and an unusual composition of PAs (epiafzelechin-derived) (Shirley *et al*., 1995; Routaboul *et al*., 2006; Schulz *et al*., 2016; Niñoles *et al*., 2023; Kerhoas *et al*., 2006). TT1, TT2, TT8, TTG1 and TT16 have a regulatory function in PA biosynthesis by activating the expression of *BANYULS* gene, encoding an anthocyanidin reductase catalysing the first committed step in PAs (Nesi *et al*., 2002; Baudry *et al*., 2004). Consequently, mutants in any of these genes are depleted in PAs but still contain flavonols (Nesi *et al*., 2002; Routaboul *et al*., 2006).

Col-0 plants had a median seed oil content of 39.2% (Figure 4b). Nearly all the mutants, either in regulatory or structural genes, displayed a significantly higher seed oil content than Col-0, from 41.1% for *tt3-7* to 52.0% for *ttg1-21* (Figure 4b), confirming previous results obtained for *tt8* and *ttg1* mutants (Chen *et al*., 2014; Chen *et al*., 2015). In contrast, we observed, as Lu *et al*. (2021), that *tt16-1* showed a small reduction in oil content (median of 39.2%) compared to Col-0 (41.7%) (Figure 4c). On the contrary, all the three *tt7* alleles showed a strong reduction in their seed oil content compared to Col-0 (26.2% for *tt7-8*, 21.6% for *tt7-9* and 25.5% for *HEM_115*). This result showed that *tt7* mutants are the only ones, among the flavonoid biosynthetic mutants studied, to display a drastic reduction in seed oil content, highlighting a specific role of *TT7* in this trait. To test whether this phenotype is linked to the role of *TT7* in the flavonoid biosynthetic pathway, double *tt7 tt4* mutants (*tt7-8 tt4-11*, *tt7-9 tt4-11* and *HEM_115 tt4-11*), which do not accumulate flavonoid were generated. As shown on Figure 4d, the seed oil content was restored to wildtype (and *tt4*) levels in all the double *tt7 tt4* mutants, suggesting that the *tt7* mutant phenotype is flavonoid-dependent.

### 6. Untargeted metabolomics reveals the metabolic signature of *tt7* mutants

To get a better picture of the seed specialized metabolites (SM, e.g. flavonoids, glucosinolates, cinnamic acids) that could be related to *tt7* phenotypes, untargeted metabolomic analyses were performed on wildtype, *tt7*, *fls1-3*, *tt3-7*, *tt4-11* simple mutants, and on *tt7 tt4* double mutants. Metabolomic analyses revealed a large diversity of SM (here named Metabolic features, Mf) with known or unknown annotation (920 Mf, Table S1). To improve the association to a known metabolic class of the metabolites with no annotation, a molecular network was built using the untargeted metabolomic data (LC-MS/MS), as in Boutet *et al*. (2022). The features with a putative annotation or metabolic category were assigned to 13 metabolic classes: flavonoids (93 Mf), glucosinolates [GSLs] (75 Mf), alkaloids (26 Mf), cinnamic acids (20 Mf), amino acids (AA) and derivatives (10 Mf), lipids and fatty acids (5 Mf), organic acids (4 Mf), phenylpropanoids (4 Mf), lignans and derivatives (3 Mf), carbohydrates (3 Mf), nucleosides and derivatives (3 Mf), terpenes (2 Mf) (Figure 5a). A principal component analysis performed using the metabolomic data separated the genotypes into three groups according to the PC1 (explaining 53% of the variance) and/or the PC2 (explaining 30% of the variance). *tt7* mutants were grouped together, highlighting a specific SM signature in the seeds of these mutants with respect to the wildtype and the other mutants in flavonoid biosynthetic genes (Figure 5b). The hierarchical cluster analysis based on SM accumulation behaviour confirmed this hypothesis, and allowed the identification of a cluster of 151 metabolic features that are highly accumulated and specific to *tt7* seeds compared to the other genotypes (Figure 5c). Among them, many known/annotated metabolites belong to the flavonoids (24), AA and derivatives (16), glucosinolates (10) and cinnamic acids (7) (Figure 5d). Flavonoids highly or exclusively accumulated in *tt7* mutants were all kaempferol decorated (glycosylated) metabolites, harbouring pentose and/or hexose sugars in their structure. These included kaempferol-rhamnose, kaempferol-rhamnose-rhamnose, kaempferol-rhamnose-pentose, kaempferol-glucose-rhamnose, and kaempferol-rhamnose-rhamnose-rhamnose (Figure 5e). Besides flavonoids, metabolomic analyses allowed the identification of other SM that were specifically accumulated in *tt7* mutants. Among them there were many AA and derivatives (mostly dipeptides), glucosinolates, including the derivative of the indole-3-carbinol GLS-degradation product and two sinapoylated-GLSs, and cinnamic acids, such as the vanillic acid, sinapoyl-pentoside, coumaroyl– and feruloyl-choline derivatives.

**Figure 5.**
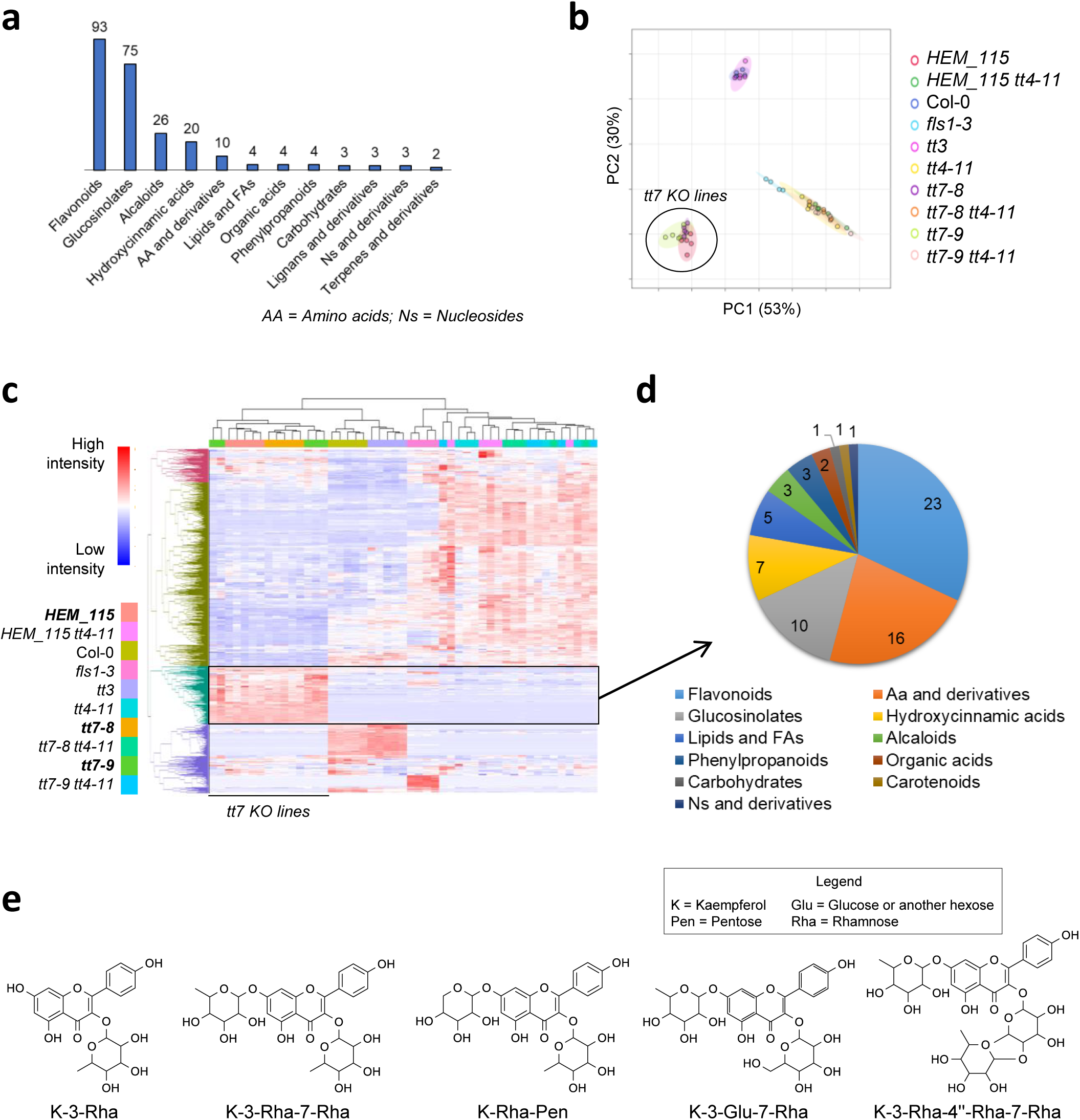
Untargeted metabolomic analyses in seeds of wildtype and flavonoid mutants. (a) Putatively annotated metabolic categories. (b) Principal component analysis of untargeted metabolomic data. Wildtype and flavonoid mutant sample distribution in seeds according to PC1 and PC2. The percentage of variance is reported for each component. (c) Hierarchical clustering and heatmap of Differentially Accumulated Metabolites (DAMs) between genotypes. (d) Annotated metabolic categories of specialized metabolites specifically accumulated in *tt7* mutants. (e) Putative structures and annotation of kaempferol-derivatives compounds highly accumulated in *tt7* mutants.

These analyses highlighted that, besides many glycosylated kaempferol derivatives, other SM were specifically accumulated in *tt7* seeds.

### 7. An over-accumulation of kaempferol-derived molecules is responsible for *tt7* seed oil phenotype

*tt7* showed a very peculiar flavonoid composition not seen in other flavonoid mutants studied, such an over-accumulation of glycosylated kaempferol flavonols and modified PA composition (data obtained in this study and in Kerhoas *et al*., 2006; Routaboul *et al*., 2006; Schulz *et al*., 2016; Niñoles *et al*., 2023). Besides kaempferol-derivatives, in this study *tt7* seeds also showed higher accumulation of many AA and derivatives, glucosinolates and cinnamic acids compared to the wild-type and the other mutants (Figure 5d). To further characterize the potential impact of kaempferol-derived molecules or the modified PA composition on *tt7* oil phenotype, *tt7 tt3* (*tt3-4 tt7cr* lines), and *tt7 fls1* (*fls1-7 tt7cr* lines) double mutants (two lines per mutant) were analysed. While *tt3-4 tt7cr* mutants accumulate kaempferol-derived molecules, but not quercetin-derived molecules nor PAs (Niñoles *et al*., 2023), *fls1-7 tt7cr* is expected to be depleted in kaempferols but would contain PAs. Seed oil content of *tt3-4 tt7cr* (Figure 4e) and *tt3-7 tt7-9* (Figure S3) mutants was similar to that of *tt7-4*. On the contrary, the two *fls1-7 tt7cr* lines showed a restored to wildtype seed oil content. These results strongly suggested that the accumulation of kaempferol-derived molecules in *tt7* was responsible for the *tt7* seed oil phenotype.

Flavonol aglycones can be glycosylated at the C3 and C7 positions, primarily with rhamnose and glucose by UDP-dependent glycosyltransferases (UGTs), including UGT78D1, UGT78D2, UGT73C6 and UGT89C1 (Saito *et al*., 2013; Tohge *et al*., 2017; Barreda *et al*., 2024, Figure 4a). UGT78D1 and UGT78D2 catalyse the transfer of rhamnose and glucose to the 3-OH position of flavonols, respectively. The major form of 7-O conjugation is 7-O-rhamnosylation, catalysed by UGT89C1. The over-accumulation of kaempferol 3-O-rhamnoside has been recently associated to other seed phenotypes observed in *tt7* mutant, including seed longevity and seed coat development (Niñoles *et al*., 2023). To determine whether the reduced seed oil content of *tt7* was also due to the accumulation of this compound, we used *tt7 ugt78d2* double mutant (Yin *et al*., 2014) as well as *ugt78d1 tt7cr* and *ugt89c1 tt7cr* lines. As shown in Figure 4f, *ugt78d1* and *ugt89c1* single mutants displayed a similar or slightly higher seed oil content than the Col-0 wildtype (39.6%, 42.5% and 40.5% respectively), whereas the *ugt78d2* mutant displayed a reduced seed oil content (34.0%). In addition, *ugt78d1 tt7cr* had a restored to wildtype seed oil content, whereas *ugt78d2 tt7-6* and both *ugt89c1 tt7cr* lines had a reduced seed oil content comparable to that of *tt7-7*. These results showed that the reduced seed oil content of *tt7* mutants is related to the over-accumulation of kaempferol 3-O-rhamnoside and potentially other glycosylated kaempferol metabolites (Figure 5e).

## DISCUSSION

### 1. Near infra-red spectroscopy combined with DSC trait allowed successful identification of mutants with impaired oil/protein correlation

Seed oil and protein contents are complex traits controlled by multiple genes and influenced by the environment and genotype × environment interaction (Li *et al*., 2006; Jasinski *et al*., 2018). Studies in oil-storing species such as rapeseed (Grami *et al*., 1977; Jolivet *et al*., 2013), sunflower (Li *et al*., 2017), soybean (Kambhampati *et al*., 2020; Chung *et al*., 2003) or the model plant *Arabidopsis thaliana* (Jasinski *et al*., 2018) have shown that the accumulation of these two components is negatively correlated. Understanding the genetic regulation of this correlation is fundamental in order to manipulate one storage compound without affecting the other. In this attempt, having in hands an accurate high-throughput phenotyping method to estimate Arabidopsis seed oil and protein contents (Jasinski *et al*., 2016), as well as a trait to estimate the strength of this correlation (DSC, Delta seed composition, Marmagne et al., 2020), we screened an EMS-mutant collection (HEM, Capilla-Perez *et al*., 2018) to identify mutants with an altered seed oil/protein relationship.

This HEM collection shows a wide range of variation for both oil (from 19.47% to 45.04%) and protein (from 14.97% to 24.70%) content. Surprisingly, this induced variation is in the same range as the natural variation we observed in Arabidopsis recombinant inbred line populations (from 23 to 48% for oil, from 14 to 29% for protein) (Jasinski *et al*., 2016). Given that seeds from the HEM collection were obtained by four successive generations of self-fertilization using a single seed descent scheme carried out in the greenhouse (i.e. under favourable growth conditions), one might have expected to observe a wider range of variation in this collection than in natural populations. Indeed, natural populations are constantly exposed to more selective environmental conditions, which can lead to many mutations being selected against. Our results suggest that oil and protein contents are constrained within these ranges and that exceeding them is either impossible or lethal. The majority of HEM lines show an increase in protein content and a decrease in oil content compared with wildtype. This is consistent with previous results obtained by screening a *Brassica napus* EMS mutant library for mutants with altered seed oil content (Tang *et al*., 2020), and suggests that it will be more difficult to positively modulate oil content than protein content.

As expected, most of the mutants follow the regression line between oil and protein content, but some deviate from the general trend (Figure 1a). This deviation is stable, and therefore genetically controlled, indicating the possibility of identifying genes controlling this deviation. To this end, a mapping-by-sequencing approach was used to identify the gene responsible for the high DSC phenotype of the *HEM_115* line. Among the nine candidate genes, *TRANSPARENT TESTA 7* (*TT7*) has a premature STOP codon, leading to a truncated protein in the *HEM_115*, making this gene the best candidate. Insertion mutant analysis as well as allelism tests and functional complementation assay demonstrate that the mutation in *TT7* is indeed responsible for the DSC phenotype of *HEM_115* (Figure 1). This shows that 1. NIRS is an accurate method for phenotyping seed oil and protein content, 2. the HEM collection is suitable for identifying mutants affected in quantitative traits such as the one studied here, 3. applying whole-genome sequencing on pool of appropriate mutant individuals from an F2 mapping population is efficient for identifying candidate genes in quantitative traits. However, the limitation of this strategy for quantitative traits comes from the phenotyping: the mutant phenotype must be sufficiently different from that of the wildtype to be able to identify individuals with a mutant phenotype in the F2 population. As an example, we applied the same approach to another *HEM* line with a phenotype close to that of Col-0 and were unable to identify a candidate region, probably due to contamination of the sequencing pool by non-mutant individuals (S. Jasinski, personal communication). The use of phenotyping robots such as the Phenoscope (Tisné *et al*., 2013) may circumvent this limitation in some cases. In this study, a large homozygous EMS mutant population was screened thanks to NIRS, allowing the detection of many mutants impaired in seed oil/protein contents and correlation. These mutants are valuable resources for studying the regulation of seed compound relative accumulation.

### 2. Seed storage compound partitioning is altered in *tt7*

*HEM_115* was strongly affected in the seed oil/protein correlation due to a significant reduction in oil compared to wildtype, with no significant variation in protein content. The other two insertion mutants in *TT7*, namely *tt7-8* and *tt7-9* gave rise to the same phenotype (Figure 1b). Lu *et al*. (2008) previously measured the carbon (C) and nitrogen (N) content of *tt7-3* seeds in order to estimate the relative abundance of seed oil and protein, respectively, and showed that the C/N ratio was decreased in *tt7-3* compared to wildtype. However, whether this decrease was due to a decrease in C, an increase in N or both was not explained in this study. An overall increase in free amino acid (FAA) content in *tt7-3* compared to the wildtype was shown, which however does not imply an increase in seed storage proteins. Indeed, FAAs, while being precursors for seed storage proteins, are also key precursors for the synthesis of several groups of primary and specialized metabolites (Amir *et al*., 2018). Here, we show that in all three *tt7* alleles studied (*HEM_115*, *tt7-8* and *tt7-9*), the oil/protein ratio decreased mainly as a result of a reduction in oil content, with no clear compensation for protein content. In addition to oil and proteins, carbohydrates are the third major component stored in mature Arabidopsis seeds and it was expected that they would increase in *tt7* mutant seeds compared to wildtype. Indeed, measurement of the total cell wall polysaccharide content of dried mature seeds showed an increase from 58 to 70% in the two *tt7* mutants analysed compared to the wildtype (Figure 3e). In Arabidopsis, which is an oleo-proteaginous species, the embryo contains mainly oil and protein, while polysaccharides are predominantly found in the seed coat, a maternal tissue. We found that the seed oil phenotype observed in *tt7* is maternally controlled (Figure 2c). Studies of other mutants affected in mucilage (*glabra2,* see also below) or flavonoid (*tt4*) biosynthesis and displaying impaired seed oil content also demonstrated that seed oil content is maternally controlled in these mutants (Shi *et al*., 2012; Xuan *et al*., 2018).

### 3. Seed coat differentiation and oil content are impaired in *tt7*

The Arabidopsis seed coat is composed of several layers of specialized tissues that accumulate a variety of metabolites (Francoz *et al*., 2018). In Arabidopsis, mucilage, a polysaccharide-rich gel, is produced by the developing seed and deposited in the outer cell layer of the seed coat. It has been shown that some mutants lacking mucilage (*transparent testa glabra1, ttg1; transparent testa 8*, *tt8*; *mucilage-modified4*, *mum4* and *glabra 2, gl2*), contain more oil in their seeds (Li *et al*., 2018; Shi *et al*., 2012; Chen *et al*., 2014), leading to the hypothesis that mucilage may compete with fatty acids (FA) for photosynthates (Shi *et al*., 2012). Indeed, sucrose, the main source of carbon, is transported from photosynthetic tissues into developing seeds where it is first unloaded into the seed coat and then imported into the endosperm to finally reach the embryo (Stadler *et al*., 2005). In the seed coat, sucrose is converted to fructose and glucose, which is then converted by the rhamnose synthase MUM4 to rhamnose, which serves as a precursor for mucilage production (Western *et al*., 2004). In the embryo, sucrose is converted by glycolysis to acetyl CoA, the key substrate for FA biosynthesis required for oil production (Baud *et al*., 2008). If sucrose is not used to produce mucilage, it can be reallocated to the FA biosynthetic pathway, resulting in seeds with increased oil content. However, disrupting the mucilage pathway downstream of the conversion of glucose to rhamnose leads to seed with wildtype oil content (*ttg2* mutants), suggesting that once glucose is converted to rhamnose, the carbon flux can no longer be redirected to FA synthesis (Shi *et al*., 2012). Mucilage production is strongly reduced in all three *tt7* alleles analysed in this study (Figure 3c), as already described for other *tt7* alleles (Niñoles *et al*., 2023). The reduced amount of mucilage could result from the starch degradation defect observed in the three *tt7* alleles studied (Figure 3b). Indeed, several studies have highlighted the accumulation of starch granules in the seed coat before and during mucilage synthesis and columella cell wall production, followed by a complete degradation of starch at seed maturity (Western *et al*., 2000; Windsor *et al*., 2000). These observations suggest that starch is necessary for either mucilage synthesis or/and columellae formation. However, studies of mutants affected either in starch metabolism such as *phosphoglucomutase 1* (*pgm1)*, or in starch degradation, such as *starch excess 1* (*sex1*) have shown that these mutants release normal amounts of mucilage (Western *et al*., 2000; Windsor *et al*., 2000). On the contrary, they display abnormal columellae, suggesting that starch may play a role as a precursor of polysaccharides used to reinforce the spindle-shaped wall in the mature cell (Western *et al*., 2000; Windsor *et al*., 2000). The observation of collapsed columella in *tt7* alleles (Figure 3d) is consistent with this hypothesis.

In addition, the defect in starch degradation observed in *tt7* could also explain the reduction in oil content. Indeed, the possibility of a metabolic competition between the starch and oil pathways has long been a matter of debate. Indeed, the biosynthesis of these two compounds depends on the availability of hexose-P in the cytosol, and mutants such as *tt7* (this study, Niñoles *et al*., 2023), *sex1* (Andriotis *et al*., 2010) or *shrunken seed1* (Lin *et al*., 2004) show a negative correlation between starch and oil content (higher starch accumulation and lower oil content). However, several counter-arguments have been put forward to refute this hypothesis. Firstly, the amount of FA is 10-fold higher than the amount of starch accumulated in the seed (Lu *et al*., 2021; Baud *et al*., 2002), suggesting that starch accumulation and subsequent degradation could not meet the requirements for oil biosynthesis. In addition, starch is still accumulated when oil synthesis and deposition have already started (Baud *et al*., 2002). Furthermore, in *B. napus*, embryo-specific downregulation of *ADP-GLUCOSE PYROPHOSPHORYLASE*, the first enzyme of the starch biosynthetic pathway, results in mature seeds with a large reduction in starch content compared to wildtype, but with a normal oil content (Vigeolas *et al*., 2004). One explanation for these conflicting results may be that in most experiments, the seed coat and embryo are examined together as a whole seed, making it impossible to distinguish the role of each compartment separately. In addition, some phenotypes may only be visible at certain stages of seed development, but not at maturity, highlighting the need for kinetic studies. In this study, we have shown that starch accumulation is specific to the seed coat (Figure 3b), but we have not measured oil content in the two compartments separately. However, given that the seed coat contains only 10% of the total seed oil, the drastic reduction in seed oil content observed in *tt7* must be due, to a large extent to a reduction in oil in the embryo.

### 4. kaempferol-3-O-rhamnoside accumulation is responsible for tt7 seed oil phenotype

The first step in flavonoid biosynthesis is the condensation of malonyl-CoA and p-coumaroyl-CoA into chalcone intermediates by the CHALCONE SYNTHASE (CHS), encoded by *TT4* (Winkel-Shirley, 2001; Saito *et al*., 2013; Lepiniec *et al*., 2006) (Figure 4a). The production of malonyl-CoA is ensured by the carboxylation of acetyl-CoA by ACETYL-COA CARBOXYLASE (ACC). It is noteworthy that malonyl-CoA is used for the synthesis of all the FA, which are then used for triacylglycerol synthesis, the main lipids found in seed oil (Rawsthorne, 2002; Ohlrogge and Jaworski, 1997; Baud *et al*., 2008). This highlights malonyl-CoA as a common source for both the seed oil and flavonoid pathways. Furthermore, many mutants affected in the flavonoid pathway such as *tt4*, *tt2*, *tt8*, and *ttg1* have been shown to have increased seed oil content compared to the corresponding wildtype (Li *et al*., 2018; Xuan *et al*., 2018; Chen *et al*., 2012; Chen *et al*., 2014; Chen *et al*., 2015), suggesting a compensation mechanism between flavonoid and oil content. However, both *tt16* (Routaboul *et al*., 2006; Lu *et al*., 2021) and *tt7* mutants (Niñoles *et al*., 2023, this study) show a concomitant reduction in flavonoid and oil contents, highlighting a more complex interplay between flavonoid and oil synthetic pathways. To gain further insight into this interaction, mutants of genes involved in flavonoid biosynthesis were studied for their oil content. A statistically significant increase in seed oil content was observed in mutants affected in regulatory genes such as *TT8* and *TTG1*, as well as in the mutant affecting the structural *TT4* gene (Figure 4b).This increase was 20, 29, and 8% in the *tt8*, *ttg1* and *tt4* mutants, respectively, which is much lower than the previously reported increases of 64, 60, and 48% in mutants affected in the same genes (Chen *et al*., 2014, 2015; Xuan *et al*., 2018). No statistically significant difference in seed oil content was observed for *tt2* in our study whereas an increase of 95% was reported for this mutant by Chen *et al*. (2012). For *ttg1* and *tt4*, these differences may be due to the fact that different alleles were studied. Furthermore, as seed oil content is strongly influenced by environmental conditions (Li *et al*., 2006; Jasinski *et al*., 2018), different results can be expected from one laboratory to another depending on the culture conditions. It is conceivable that a mutation altering the pathway prior to the conversion of dihydrokaemfperol to dihydroquercetin by TT7 could result in a redirection of malonyl-CoA towards FA biosynthesis, but the fact that mutants affected in genes both upstream (*tt4*, *tt6*) and downstream (*tt2*, *tt8*, *ttg1*, *tt11*) of *TT7* showed an increase in oil content rules out this hypothesis.

Of all the flavonoid mutants examined in this study, only *tt16* and *tt7* showed a reduction in seed oil content compared to the wildtype, with *tt7* showing the most dramatic reduction. Consistent with this phenotype, *tt7-8* had a reduced level of expression of *WRINKLED1* (*WRI1*), a gene encoding a key transcription factor in the regulation of plant oil biosynthesis (Focks and Benning, 1998; Baud *et al*., 2007; Cernac and Benning, 2004), and two of its targets: *PLASTIDIAL PYRUVATE KINASE β1* (*PKp-β1*) and *BIOTIN CARBOXYL CARRIER PROTEIN 2* (*BCCP2*) (Baud *et al*., 2009; Maeo *et al*., 2009). A downregulation of *PKp-β1* and *BCCP2* was also reported in a transcriptomic analysis of *tt7-7* seeds (Niñoles *et al*., 2023). However, *WRI1* was not identified as a differentially expressed gene (DEG) in this study, perhaps because of its very low expression levels. Notably, the transcriptomic analysis of Niñoles *et al*. (2023) revealed a downregulation of genes encoding several lipid metabolism enzymes in the *tt7-7* mutant, among other metabolic processes. Taken together, our results suggest that *TT7* positively acts on the oil biosynthetic pathway, at least via the activation of *WRI1*, by a mechanism that is still unknown.

*tt7* mutants have a very specific flavonoid composition not seen in other flavonoid mutants (Routaboul *et al*., 2006, this study), in particular a strong accumulation of kaempferol glycosides and no quercetin-derived glycosides, suggesting that the reduction in seed oil content may be related to this unique flavonoid composition. In wildtype Arabidopsis seeds, UGT78D1, UGT78D2 and UGT89C1 are the main enzymes involved in flavonol glycosylation (Tohge *et al*., 2017; Saito *et al*., 2013). We show that *ugt78d2* and *ugt89c1* mutations in combination with *tt7* do not allow recovery of the wildtype phenotype, whereas *ugt78d1* mutation is able to abolish the *tt7* seed oil phenotype (Figure 4f), highlighting a role for kaempferol-3-O-rhamnoside in the *tt7* phenotype. In wildtype seeds, kaempferol-3-O-rhamnoside is mainly found in the seed coat (Routaboul *et al*., 2006), probably due to the expression profile of *UGT78D1*, which is restricted to the seed coat at the beginning of seed development (Belmonte *et al*., 2013). The expression profile of *UGT89C1*, encoding F7RHAT, which performs the next step of glycosylation, is later in seed development, and both in the embryo and in the seed coat (Belmonte *et al*., 2013). The seed coat-specific localisation of kaempferol-3-O-rhamnoside is consistent with the maternal control of seed oil content by *TT7*. Interestingly, the *ugt78d2* single mutant, in which flavonol 3-O-glycosylation may switch from 3-O-glucoside to 3-O-rhamnoside via UGT78D1, and then possibly increase the levels of kaempferol-3-O-rhamnoside and quercetin-3-O-rhamnoside, has a reduced seed oil content, although not as much as *tt7*. This could be explained by a moderate increase in kaempferol-3-O-rhamnoside in *ugt78d2* compared to *tt7*, suggesting a negative correlation between kaempferol-3-O-rhamnoside levels and seed oil content.

### 5. Role of flavonols in plant development

Flavonols play many roles in plant development and in recent decades more and more specific functions have been assigned to certain compounds, mainly through the use of mutants. In this study, we show for the first time that an increase in kaempferol-3-O-rhamnoside levels in *tt7* results in a drastic reduction in seed oil content, a phenotype never described for this mutant until now. In addition, we confirm that this increase in kaempferol-3-O-rhamnoside levels is responsible of impaired starch degradation and mucilage defects in *tt7* as already shown by Niñoles *et al*. (2023). Additional phenotypes of *tt7*, such as suberin biosynthesis and seed longevity were described by Niñoles *et al*. (2023). Taken together, these results indicate that kaempferol-3-O-rhamnoside plays an important role in seed coat differentiation, seed oil content and seed physiology. Previously, Ringli *et al*. (2008), show that a large increase in kaempferol-3-O-glycoside and another unknown kaempferol-derived compound is probably responsible for the *repressor of lrx1-2* (*rol1-2*) mutant phenotype, including hyponastic growth, aberrant pavement cell and stomatal morphology in cotyledons, and defective trichome formation. The *male sterility 1* (*ms1*) mutant, which is defective in normal pollen and tapetum development, is depleted in two flavonol derivatives (kaempferol 3-O-glucosyl-(1→2)-glucoside and quercetin 3-O-glucosyl-(1→2)-glucoside) (Yonekura-Sakakibara *et al*., 2014). Mutants in the *UGT78D2* gene exhibit a dwarf stature and increased branching, as well as a reduced polar auxin transport (PAT) in the shoot, phenotypes that have been attributed to increased levels of kaempferol 3-O-rhamnoside-7-O-rhamnoside (Yin *et al*., 2014). Furthermore, by analysing auxin transport in several genotypes with different levels of kaempferol 3-O-rhamnoside-7-O-rhamnoside, Yin *et al*. (2014) observed an inverse correlation between basipetal auxin transport and kaempferol 3-O-rhamnoside-7-O-rhamnoside levels.

In addition, several experiments with *tt* mutants have shown that auxin transport is enhanced in the absence of flavonoids (in the *tt4* mutant) and reduced in the presence of excess flavonols (in the *tt7* and *tt3* mutants) (Buer and Muday, 2004; Peer *et al*., 2004; Kuhn *et al*., 2011; Peer and Murphy, 2007). Furthermore, it has been shown that flavonols can compete with the auxin transport inhibitor 1-N-naphthylphthalamic acid (NPA) for binding to proteins involved in auxin transport (Murphy *et al*., 2000). Recently, to gain insight into the effects of flavonols on global gene expression and signalling pathways, Naik *et al*., (2024) performed comparative transcriptome deep sequencing (RNA-seq) and targeted metabolite profiling of the flavonol-deficient *fls1* mutant following exogenous treatment with different flavonols. Their transcriptome analysis revealed several DEGs in the flavonoid biosynthetic pathway itself but also in other pathways, such as phytohormone signalling, stress responses, suberin biosynthesis, and specialized metabolism, highlighting the role of flavonols in transcriptional regulation of various developmental and signalling processes. Interestingly, Grunewald *et al*., (2012) have shown that both *TT4* and *TT7* transcripts are significantly up-regulated in response to an auxin treatment in roots, and that this regulation occurs via the transcription factor WRKY23. These results highlight a possible feedback loop between flavonols regulating auxin accumulation by acting on PAT, and auxin regulating flavonol synthesis via transcriptional regulation of *TT* genes. Recently, Kong *et al*., (2017) have unravelled a link between auxin and the oil biosynthetic pathway via the role of the protein WRI1. They showed that AtWRI1 binds *in vitro* to the promoter of *GRETCHEN HAGEN3.3* (*GH3.3*), which encodes an enzyme that converts active IAA (indole-3-acetic acid) to inactive amide-IAA conjugates, and to the promoters of *PIN-FORMED*4 and *5* (*PIN4* and *PIN5*). In agreement with these data, they observed a higher level of the IAA-Asp conjugate and a decrease in PAT in *wri1* mutant seedlings, suggesting that WRI1 may acts positively on PAT. Taken together, these results suggest complex regulatory mechanisms between auxin, flavonols and oil biosynthetic pathways. However, most of these experiments have been carried out on roots or seedlings, and whether these regulations take place in the seed needs further investigation. Our results pave the way to further investigate the interplay between these three pathways in the seed.

## EXPERIMENTAL PROCEDURES

### 1. Plant materials and growth conditions

HEM lines were obtained from the Versailles Biological Resource Centre (https://publiclines.versailles.inrae.fr/). T-DNA insertion mutants *tt7-7* (GK_629C11) (Appelhagen *et al*., 2014), *tt7-8* (SALK_039417), *tt7-9* (SALK_seq_064306), *tt2-5* (SALK_005260), *tt8-7* (SALK_063334), *ttg1-21* (GK-580A05), *tt4-11* (SALK_020583) (Buer *et al*., 2006), *tt1-3* (SALK_026171), *tt6-3* (SALK_113904) (Owens, Crosby, *et al*., 2008), *ugt78d1* (SAIL_568_F08), *ugt78d2* (SALK_049338), *ugt89c1* (Salk_seq_068559.0) were ordered from the Nottingham Arabidopsis Stock Centre (http://arabidopsis.info). *fls1-3* (Kuhn *et al*., 2011) and *tt3-7* (SALK_099848C) (Chapman and Muday, 2021) were kindly provided by G. Muday. *tt3-4*, *tt7-4* (Routaboul *et al*., 2006) and *fls1-7* (FLAG-533E06) (Owens, Alerding, *et al*., 2008) were kindly provided by I. Debeaujon. *tt11-11* (SALK_073183) (Bowerman *et al*., 2012) was kindly given by B. Winkel. *tt16-1* (Nesi *et al*., 2002) was kindly provided by E. Magnani. *ugt78d2 tt7* (Yin *et al*., 2014) was obtained from Anthony Schäffner. The mutation in *tt7* is a deletion of 10 bp in the first exon, that results in a truncated protein. *tt3-4 tt7cr* (L8 and L20), *fls1-7 tt7cr* (L38 and L47), *ugt78d1 tt7cr* (L126) and *ugt89c1 tt7cr* (L25 and L31) correspond to the indicated mutant CRISP-edited for *TT7* and are described in Niñoles *et al*. (2023). The *tt7-8 tt4-11*, *tt7-9 tt4-11*, *HEM_115 tt4-11* and *tt3-7 tt7-9* double mutant lines were generated by crossing the corresponding alleles. All mutants (except *tt7-4* and *tt3-4*) were confirmed by PCR genotyping using specific primers listed in Table S2. For the Salk mutants, the Salk T-DNA specific primer LB-Salk2 (5’-GCTTTCTTCCCTTCCTTTCTC-3’) was used, except for the *tt11-11* and *ugt89c1*, which were genotyped with the LBb1.3 primer (5’-ATTTTGCCGATTTCGGAAC-3’). For the Gabi-Kat mutants, the Gabi T-DNA specific primer Gabi_o8409 (5’-ATATTGACCATCATACTCATTGC –3’) was used. For the FLAG mutants, the specific primer FLAG_LB4 (5’-CGTGTGCCAGGTGCCCACGGAATAGT-3’) was used. For the SAIL mutants, the specific primer Sail_LB1 (5’-GCCTTTTCAGAAATGGATAAATAGCCTTGCTTCC-3’) was used. The T-DNA specific primers were used with the reverse specific primers to detect the insertion. All plants were grown on soil in greenhouse under natural light supplemented with sodium lamps to provide a 16 h photoperiod at 22°C. Depending on the background of each mutant, the Columbia-0 (Col-0) or the Wassilewskija-0 (Ws-0) ecotype was used as the wildtype control. In all the analyses, comparisons were made between seed lots from plants that had been grown and harvested simultaneously.

### 2. Whole Genome Sequencing and Mutation Analysis

Twenty-four F3 lines from the *Col-0* x *HEM_115* cross, that was supposed to carry the mutation of interest at the homozygous state (see Results section), were chosen in order to extract their genomic DNA for deep sequencing in pool. F3 seeds were sown on agar plates with Arabidopsis media. After 14 days, plantlets were sampled and their genomic DNA was extracted using CTAB (Cetyl trimethyl ammonium bromide) buffer. Genome sequencing was performed with Illumina Hiseq3000 with a 32X coverage.

The WGS reads were cleaned up using the Trimmomatic tool (0.32) (Bolger *et al*., 2014), and then mapped to A. thaliana TAIR10 with BWA mem (0.6.1) software (Li and Durbin, 2009). Afterwards alignment files were processed according to the genome analysis toolkit (GATK) Best Practices recommendations (Depristo *et al*., 2011). SNVs (single nucleotide variations) were called using GATK (3.5) HaplotypeCaller algorithm (Poplin *et al*., 2017) across all samples simultaneously, and annotated using SnpEff tools (3.6c) (Cingolani *et al*., 2012). To specifically select for EMS-induced mutation, we removed false positives and polymorphisms (distance between the studied variety and the reference genome). For this purpose, a filtering process was applied based on quality and coverage (QUAL > 80, total depth > 8, reads supporting the variant allele > 1). Additionally, we kept only substitutions that correspond to typical EMS-induced mutations (G>A and C>T) and filtered out SNVs that were common across the sample set. The frequency of the alternate allele relative to reference genome was visualized across the genome (position) to identify the peak that likely corresponds to interval containing the mutation(s) responsible for the phenotype of interest. These mutations were further selected based on annotation criteria (within genes, predicted impact of the modifications…).

### 3. Plasmid construction and plant transformation

All vectors used in this work have been designed using GoldenBraid system following the described assembly strategy (Sarrion-Perdigones *et al*., 2013). The pTT7::TT7::Tnos construct was assembled into pDGB3_alpha1 (#68228 from Addgene) using the three modules pTT7, TT7, tNos (GB0037) domesticated into pUPD2 (#68161 from Addgene). pTT7 and TT7 modules were domesticated from genomic DNA using the GB Domesticator tool (https://gbcloning.upv.es/do/domestication/). TT7 coding sequence displays a *BsaI* restriction site and was cloned in two fragments. The pCMV::DsRed::tNos construct into pDGB3_alpha2 (#68229 from Addgene) was a gift from Z. Kelemen (IJPB). The two transcriptional units were then assembled into pDGB3_omega1 (#68238 from Addgene). All constructions were checked by sequencing of the entire module for domestication or the junctions between the different modules in alpha vectors. The presence of the DsRed selection marker was checked using appropriate primers. All primers used for domestication and sequencing are provided in Table S3.

E. coli strain NEB10 beta electromax (NEB) was used as bacterial host. Transformed bacterial clones were isolated on LB agar plates containing 50 µg/ml chloramphenicol for the pUPD2 backbone, 100 µg/ml kanamycin for alpha-level backbones and 50 µg/ml spectinomycin for omega-level backbones. *Agrobacterium tumefaciens* strain C58C1 (pMP90) was used as bacterial host for plant transformation. LB agar selection plates containing 50 µg/ml rifampicin, 25 µg/ml gentamicin and 50 µg/ml spectinomycin were used to isolate transformed bacterial clones.

*Arabidopsis thaliana* (Col-0, *HEM_115* or *tt7-8*) were transformed by floral dipping (Bechtold and Pelletier, 1998), with a minor modification: 2 min dipping into a solution (sucrose 5% + 0.2 ml Silwet-77/L) containing *A. tumefaciens* strain C58C1 (pMP90). Fluorescent seeds, containing the DsRed, were identified under a Leica DMS1000 microscope with DsRed filter. Fluorescent T1 seeds were selected and put on soil to obtain the next T2 generation. The T2 seed populations contained three types of seeds: non-fluorescent (not transformed), moderately fluorescent (hemizygous for the T-DNA) and strongly fluorescent (homozygous for the T-DNA) seeds. Counting for the three types of seeds allowed identifying lines carrying insertion at one locus. Strongly fluorescent seeds from these lines were grown to obtain homozygous seeds of the third (T3) generation. The NIRS analyses were carried out on seeds from the T3 or the following generation.

### 4. Expression analysis by RT-qPCR

For RT-qPCR analysis, Col-0 and *tt7-8* plants were grown in greenhouse under long day condition. The time of pollination (0 d after pollination [DAP]) was defined phenotypically as the time at which the flowers are just starting to open and the long stamens grow over the gynoecium. Every two days, during ten days, flowers at this stage were tagged with coloured thread. The colour of the thread identified the date of pollination and allowed the selection of developing siliques at precise ages. Four siliques from 4 different plants were then dissected and seeds were pooled for total RNA extraction using the RNeasy Plant Mini Kit (Qiagen) following the manufacturer’s protocol. DNAse treatment was performed on columns. Two hundred and fifty nanograms of total RNA was reverse transcribed by the RevertAid M-MuLV Reverse Transcriptase (Fermentas) with an oligo (dT) primer according to the manufacturer’s protocol. cDNA were diluted 20 times and 4.4 µL were used as template in a 20 µL qPCR reaction. PCR primers specific for *WRI1*, *BCCP2*, *PKp-β1*, *EF1a* and *PP2AA3* were used and all sequence primers are described in Table S4. *EF1a* and *PP2AA3* were used as reference genes.

### 5. Biochemical and physiological analysis of seeds

#### Seed oil content determination

Seed oil content was estimated by NIRS as described in Jasinski *et al*. (2016). Total seed fatty acid content and fatty acid composition were determined by direct transmethylation followed by gas chromatography as described in Jasinski *et al*. (2012).

#### Mucilage staining

Ruthenium red (Sigma) was dissolved in water at a concentration of 0.02% (w/v). Seeds were shacked 10 min in water and non-adherent mucilage was removed by pipetting. Seeds were gently pipetted on a slide and adherent mucilage was stained directly by adding the 0.02% Ruthenium red solution for 10 minutes and observed with a light microscope (Axioplan 2; Zeiss).

#### Starch staining

Mature seeds were placed on Whatman paper with water at 4°C for 2h to facilitate the dissection. Embryos and seed coats were separately placed in 70% (v/v) ethanol and boiled for 5 min, then incubated in Lugol solution (Sigma, ref 32922) for 20 min. Excess staining solution was removed by washing with distilled water.

#### Clearing Seeds and Imaging

For embryo developmental analysis, flowers at the time of pollination (0 day after pollination [DAP]) were marked with different colour threads every two days during 12 days, allowing the selection of silique at precise ages. Siliques were dissected and seeds were mounted in clearing solution (chloral hydrate:glycerol:water (8:1:2, w:v:v). After few hours, slides were observed by differential interference contrast microscopy with an Axioplan 2 microscope (Zeiss).

#### Scanning electron microscopy

Dry mature seeds were mounted onto a Peltier cooling stage with adhesive discs (Deben) and observed with a SH-1500 tabletop scanning electron microscope (Hirox).

#### Polysaccharides content

Dry mature seeds (100 mg) were dried and then milled using FastPrep-24 instrument (MP Biochemicals, CA) at a speed of 6.5 m/s for 60 s. Each mill was then defatted with 1 mL hexane during 1 hour at 40°C and the defatted residue was recovered after centrifugation (30 min, 15 000 *g*). Defatted mills were dried before Alcohol Insoluble Materials (AIMs) extraction using the ASE® 350 (Thermo Scientific Dionex) as previously described with minor modifications (Le Gall *et al*., 2023). Briefly, approximately 60 mg of defatted mills were weighted and AIMs were extracted during 10 min at 100°C. The osidic composition of cell wall polysaccharides were then performed as described in Le Gall *et al*. (2023).

#### Seed area

Pictures of seeds were taken with an Axiocam 208 color camera (Zeiss) attached to a MZ6 microscope (Leica). Seed area was determined using ImageJ software (https://imagej.nih.gov/ij/index.html).

### 6. Statistical analyses

All statistical analyses were performed using the free software environment R Version 4.2.1 (https://www.r-project.org/).

### 7. Seed specialized profiling and annotation using UPLC–MS/MS untargeted metabolomics

Untargeted metabolomic analyses were carried out following the protocol described in Boutet *et al*. (2022), with modifications. Polar and semi-polar metabolites were extracted from 10 mg of *A. thaliana* mature dry seeds. For metabolite extraction, 1 ml of methanol/acetone/water/trifluoroacetic acid (30/42/28/0.1 v/v/v/v) extraction buffer, conserved at 4°C, and 500 ng of Apigenin (used as internal standard) were added to each sample, which was then homogenized in 2-ml tubes using a FastPrep instrument (1 min, 14 000 rpm). The mixtures were then shaken for 30 min at 4°C using a ThermoMixerTM C (Eppendorf), placed in an ice-cooled ultrasonication bath for 2 min, and centrifuged at 10000 rpm for 10 minutes (4°C). The supernatant was collected in 4 ml glass tube. 1 ml of extraction buffer was added to the pellet and the extraction procedure was repeated. The supernatant was added to the glass tube containing the previously extracted mixture. The mixture containing the polar/semi-polar metabolites was dried down in a SpeedVac vacuum concentrator (o/n) and resuspended in 200 μl H20 + 10% acetonitrile just before injection.

Metabolomic data were acquired using a UHPLC system (Ultimate 3000 Thermo) coupled to quadrupole time of flight mass spectrometer (Q-Tof Impact II Bruker Daltonics, Bremen, Germany) using the protocol described in Boutet *et al*. (2022).

The raw data files (.d; Bruker Daltonics, Bremen, Germany) were converted to .mzXML format using the MSConvert software (ProteoWizard package 3.0). mzXML data processing, mass detection, chromatogram building, deconvolution, samples alignment and data export were performed using MZmine 3.0 software (http://mzmine.github.io/) for both positive and negative data using the same parameters indicated in Boutet *et al*. (2022). Metabolic features intensity was normalized according to the internal standard (Apigenin, 500 ng) and weight of seeds used for the extraction. A first research in the library with Mzmine was carried out.

The obtained RT and m/z data of each feature were initially compared with the IJPB homemade library containing more than 200 standards or experimental common features (RT, m/z). Molecular networks were then generated with MetGem software (Olivon *et al*. (2018); https://metgem.github.io) using the .mgf and .csv files obtained with MZmine3.0 analysis. ESI+ and ESI-molecular networks were generated using cosine score thresholds of 0.75 and 0.8, respectively. Molecular networks were exported to Cytoscape software (Shannon *et al*. (2003); https://cytoscape.org/) to format the metabolic categories. Next, the metabolomic data used for molecular network analyses were searched against the available MS2 spectral libraries (Massbank NA, GNPS Public Spectral Library, NIST14 Tandem, NIH Natural Product and MS-Dial), with absolute m/z tolerance of 0.02, 4 minimum matched peaks and minimal cosine score of 0.8. Not-annotated metabolites that belong to molecular network clusters containing annotated metabolites from the molecular network analysis were assigned to the same chemical family. Finally, the putative structure and assignment to a metabolic category of selected metabolic features that had no or unclear annotation was performed with Sirius software (https://bio. informatik.uni-jena.de/software/sirius/).

## 8. Statistical and clustering analyses of untargeted metabolomic data

Statistical analyses to perform a principal cluster analysis (PCA) and identify Differentially Accumulated Metabolites (DAMs) were performed using MetaboAnalyst 5.0 (Pang *et al*., 2022; Xia *et al*., 2009). In particular, one-way Analysis of Variance (ANOVA) was performed to identify the metabolites showing statistically different accumulation among the genotypes (i.e. wildtype and knockout mutants, adjusted p value 0.05, Fisher’s LSD). Hierarchical clustering according to SM accumulation behaviour was performed using *pheatmap* R package.

## Acknowledgments

We are very grateful to the IJPB’s Plant Observatory platforms Versailles Arabidopsis Stock Center (PO-VASC, https://publiclines.versailles.inrae.fr/) for giving us access to the seeds from the HEM library. We thank Dr Isabelle Debeaujon, Dr E. Magnani, Dr G. Muday, Dr A. Schäffner and Pr B. Winkel for kindly sharing Arabidopsis mutant lines. We thank N. Bessoltane for her help with sequence analysis. We thank S. Baud, S. Bonhomme, B. Dubreucq, R. Le-hir, L. Lepiniec, H. North, and A. de Saint Germain for fruitful discussions. We thank Lilian Dahuron and Philippe Marechal for plant care. We thank Lucie Le Bot for cell wall polysaccharide analyses. Cell wall analyses were performed on the BIBS instrumental platform (DOI: 10.15454/1.5572358121569739E12; http://www.bibs.inrae.fr/bibs_eng/, UR1268 BIA, IBiSA, Phenome-Emphasis-FR ANR-11-INBS-0012, PROBE infrastructure, Biogenouest). This work has benefited from the support of IJPB’s Plant Observatory platforms: the plant growth observatory platform (PO-Plants) and the Chemistry and Metabolism plateform (PO-Chem). The IJPB benefits from the support of Saclay Plant Sciences-SPS (ANR-17-EUR-0007). IJPB and BIA experimentations were supported by Promosol.

## Author contributions

S.J. and P.G. conceived and supervised the study. M.C. and S.B. designed and performed metabolomic analysis, analysed the data and M.C. wrote the corresponding sections. S.L.G. designed and performed cell wall analysis and wrote the corresponding sections. R.N. and J.G. made the CRISPR-edited *tt7* lines. A.L., P.G. and S.J. performed all the other experiments and S.J. and P.G. analysed the data. S.J. wrote the manuscript. All authors read and approved the manuscript.

## Availability of data and material (data transparency)

The sequencing raw data of the *HEM_115* line is available in ArrayExpress (E-MTAB-14527). The metabolomic data and metadata have been deposited in MassiVE (doi:10.25345/C5X05XQ87).

## Supporting Information

**Figure S1**. Mapping-by-sequencing of the *HEM_115* mutant leads to the identification of *TT7* (a) Median of seed oil and protein content of eleven *HEM* mutants and the wildtype are represented. Col-0 is indicated in black, *HEM* lines are in dark grey. Bars show the 25–75% quartiles (6 ≤ n ≤ 10 for all genotypes). No bar is indicated for mutant *HEM_026* for which only one plant gave enough seeds. A Kruskal-Wallis statistical test, followed by a post-hoc Man & Whitney test were performed to compare each mutant to Col-0 for both traits. Significance of comparison is indicated in bracket (oil/Protein), ***P < 0.001, **P < 0.01 and *P < 0.05, ns=not significant.

(b, c) Scatter plots of NIRS versus chemical method values for oil % (b) and protein % (c) for nine genotypes. The regression line is indicated in black and the confidence interval in grey. The Pearson correlation coefficient R and the p value are indicated on the top of each graph.

(d) 150 F3 seed lots from the Col-0 x *HEM_115* cross (grey dots and triangles) were phenotyped by NIRS as well as seeds from Col-0 (black dots) and *HEM_115* (red dots). F3 seed lots displaying a mutant phenotype were chosen for deep sequencing (grey triangles).

(e) amino-acid alignment of TT7 protein of Col-0 and *HEM_115*.

(f) fatty acid composition of mature seeds of Col-0, *HEM_115*, *tt7-8* and *tt7-9*. 100 seeds from 4 plants cultivated in the same time were analysed for each genotype. A Kruskal-Wallis statistical test, followed by a post-hoc Man & Whitney test were performed to compare each mutant to Col-0 for each fatty acid. Significance of comparison is indicated, **P < 0.01.

**Figure S2**. All cell wall saccharides are increased in *tt7* mutants compared to the wildtype.

Median of rhamnose (a), galacturonic acid (b), mannose (c), glucose (d), xylose €, galactose (e), and arabinose (f) in Col-0, HEM_115 and tt7-8. Boxes show the 25–75% quartiles, the median value (inner horizontal line), and whiskers extending from the lower to the upper adjacent values. A Kruskal-Wallis statistical test, followed by a post-hoc Man & Whitney test were performed to compare each mutant to Col-0. Significance of comparison is indicated, *P < 0.05, **P < 0.01.

**Figure S3**. The *tt3 tt7* double mutant shows the same seed oil phenotype than *tt7*.

Median seed oil content of *tt7-9*, *tt3-7* and the *tt3-7 tt7-9* double mutant. Boxes show the 25–75% quartiles, the median value (inner horizontal line), and whiskers extending from the lower to the upper adjacent values. A Kruskal-Wallis statistical test, followed by a post-hoc pairwise Wilcoxon test were performed to compare all pairs of genotypes. Genotypes with the same letter are not significantly different from each other, while genotypes with different letters are significantly different (P ≤ 0.05).

