## Supplemental figures for "Innovative mutant screening identifies *TRANSPARENT TESTA7* as a player in seed oil/protein partitioning"

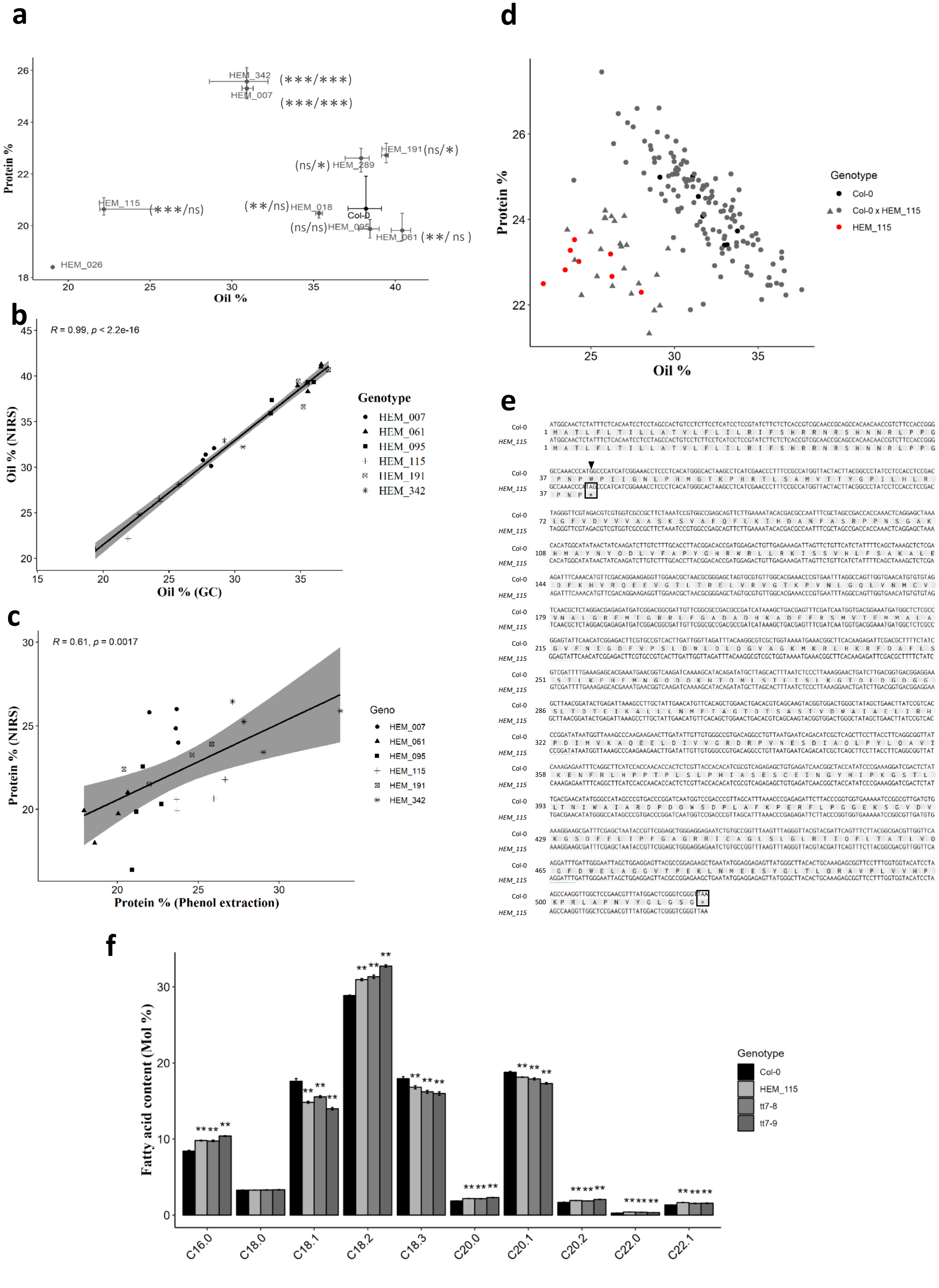


**Supplemental Figure S1**

**Figure S1.** Mapping-by-sequencing of the *HEM_115* mutant leads to the identification of *TT7*

(a) Median of seed oil and protein content of eleven *HEM* mutants and the wildtype are represented. Col-0 is indicated in black, *HEM* lines are in dark grey. Bars show the 25–75% quartiles ($6 \leq n \leq 10$ for all genotypes). No bar is indicated for mutant *HEM_026* for which only one plant gave enough seeds. A Kruskal-Wallis statistical test, followed by a post-hoc Man & Whitney test were performed to compare each mutant to Col-0 for both traits. Significance of comparison is indicated in bracket (oil/Protein), ***$P < 0.001$, **$P < 0.01$ and *$P < 0.05$, ns=not significant.

(b, c) Scatter plots of NIRS versus chemical method values for oil % (b) and protein % (c) for nine genotypes. The regression line is indicated in black and the confidence interval in grey. The Pearson correlation coefficient R and the p value are indicated on the top of each graph.

(d) 150 F3 seed lots from the Col-0 x *HEM_115* cross (grey dots and triangles) were phenotyped by NIRS as well as seeds from Col-0 (black dots) and *HEM_115* (red dots). F3 seed lots displaying a mutant phenotype were chosen for deep sequencing (grey triangles).

(e) amino-acid alignment of TT7 protein of Col-0 and *HEM_115*.

(f) fatty acid composition of mature seeds of Col-0, *HEM_115*, *tt7-8* and *tt7-9*. 100 seeds from 4 plants cultivated in the same time were analysed for each genotype. A Kruskal-Wallis statistical test, followed by a post-hoc Man & Whitney test were performed to compare each mutant to Col-0 for each fatty acid. Significance of comparison is indicated, **$P < 0.01$.

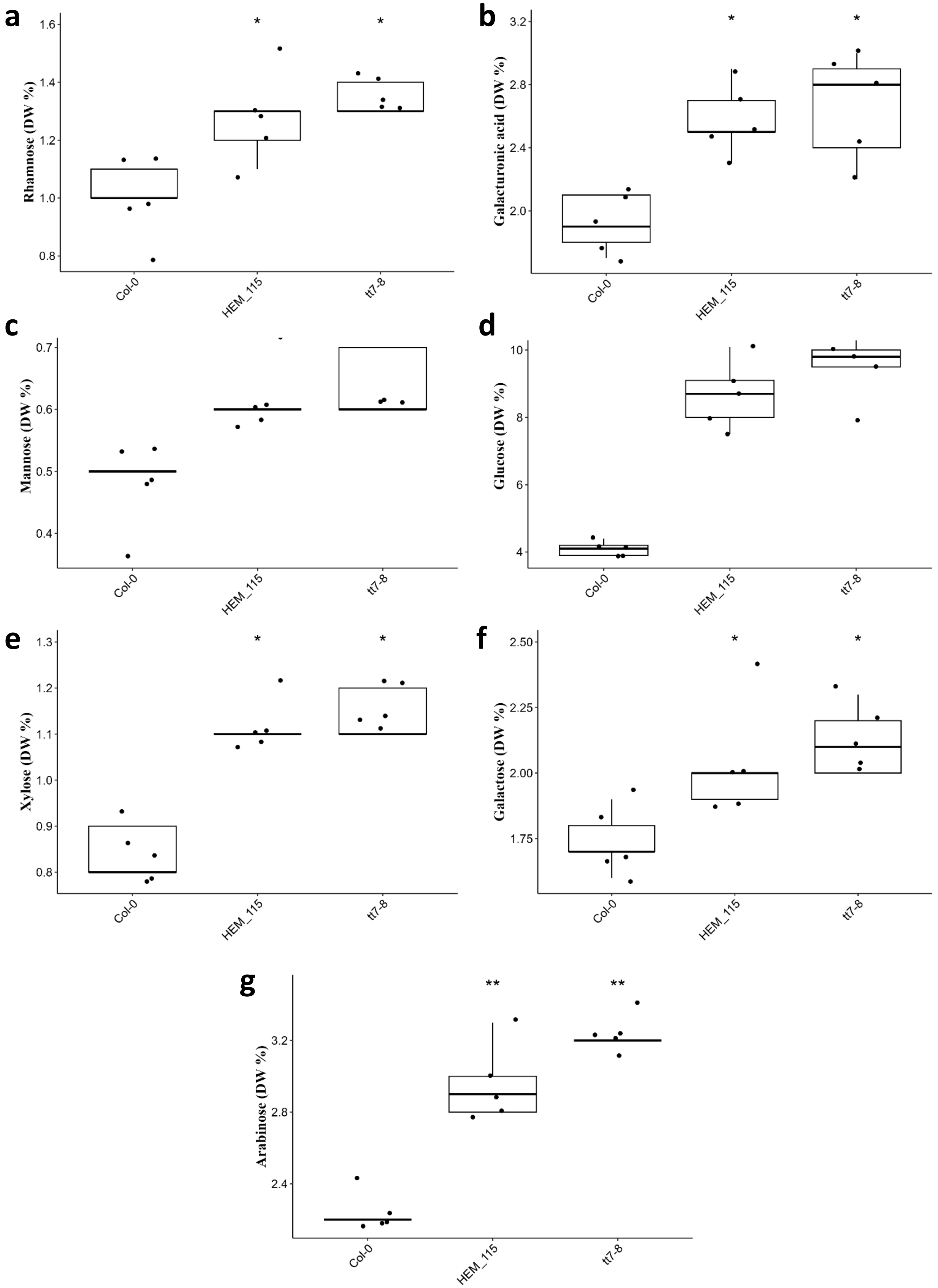


**Supplemental Figure S2**

**Figure S2.** All cell wall saccharides are increased in *tt7* mutants compared to the wildtype. Median of rhamnose (a), galacturonic acid (b), mannose (c), glucose (d), xylose €, galactose (e), and arabinose (f) in Col-0, HEM_115 and tt7-8. Boxes show the 25–75% quartiles, the median value (inner horizontal line), and whiskers extending from the lower to the upper adjacent values. A Kruskal-Wallis statistical test, followed by a post-hoc Man & Whitney test were performed to compare each mutant to Col-0. Significance of comparison is indicated, *P < 0.05, **P < 0.01.

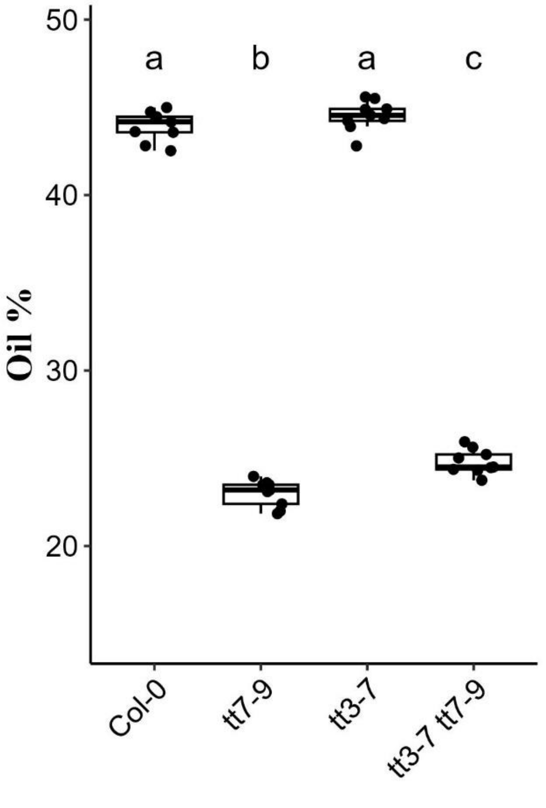


**Supplemental Figure S3**

**Figure S3.** The *tt3 tt7* double mutant shows the same seed oil phenotype than *tt7*.
Median seed oil content of *tt7-9*, *tt3-7* and the *tt3-7 tt7-9* double mutant. Boxes show the 25–75% quartiles, the median value (inner horizontal line), and whiskers extending from the lower to the upper adjacent values. A Kruskal-Wallis statistical test, followed by a post-hoc pairwise Wilcoxon test were performed to compare all pairs of genotypes. Genotypes with the same letter are not significantly different from each other, while genotypes with different letters are significantly different ($P \leq 0.05$).
